# Structural generalization and continual learning enabled by factorized entorhinal-hippocampal memory and entorhinal-parietal action circuits

**DOI:** 10.64898/2026.08.25.747129

**Authors:** Jaedong Hwang, Sujaya Neupane, Mehrdad Jazayeri, Ila Fiete

**Affiliations:** Department of Brain & Cognitive Sciences, McGovern Institute for Brain Research, MIT; Department of Electrical Engineering and Computer Science, MIT; Coherent Research, Montreal, Canada; Howard Hughes Medical Institute

## Abstract

Flexible behavior requires generalizable memory and learning. For example, we rapidly learn to commute in new cities by reusing our knowledge of Euclidean two-dimensional space and structures like roundabouts and subway systems without forgetting how to get to a favorite restaurant back home. Yet we lack a detailed understanding of how the brain uses existing knowledge to generalize while retaining the memory of specific past experiences. To address this gap, we combine behavioral measurements, neural recordings, and computational modeling in an abstract sequential image navigation task to study three forms of generalization: mnemonic generalization, from visual to mental navigation; transitive generalization, from trained to novel routes; and structural generalization, from familiar to new environments. In contrast to monkeys and humans, recurrent neural networks failed at all generalizations. We found that a structured entorhinal-hippocampal memory model, which provides a content-independent metric scaffold based on grid cells for storing experience, coupled to a policy recurrent network, succeeds at all three. The content-independent scaffold enables mnemonic and transitive generalization through path integration and facilitates structural generalization by allowing reuse of a previously learned action policy network. Moreover, the scaffold’s high combinatorial capacity permits continual learning without catastrophic forgetting. We recorded neural activity from the entorhinal cortex and posterior parietal cortex of two monkeys performing the task and found two distinct computations across the neural population. Modularizing an entorhinal and parietal action policy network to separately track distance and initiate actions captured the distinct population dynamics and improved model performance. Finally, we added a reinforcement learning module to the network that enabled it to learn an appropriate scale factor to align the grid periodicity with the environmental temporal structure. Our findings reveal that an architecture which factorizes invariant metric representations from rapid sensory associations and a transferable policy learns, generalizes, and remembers like the brain.

## Introduction

The hippocampal formation (HF) is responsible for the formation of episodic memories (Scoville and Milner 1957) and the representation of physical space in the form of spatially tuned neural responses like place and grid cells (O’Keefe and Nadel 1978; O’Keefe and Dostrovsky 1971; Sargolini et al. 2006; Hafting et al. 2005). Episodic memory formation has been hypothesized to generate a foundation for flexible mnemonic behavior by constructing an auto-associative memory buffer for events and episodes (Treves and Rolls 1994) that can be referenced in the future. The spatial representations in the hippocampus have been proposed to take the form of spatial *cognitive maps* which associate sensory experiences with coordinate estimates generated from self-movement (Tolman 1948; O’Keefe and Nadel 1978; Whittington et al. 2022). The idea of spatial cognitive maps is closely related to the computational framework of simultaneous localization and mapping in robotics, with striking parallels between the two fields (Widloski and Fiete 2014; Kanitscheider and Fiete 2017; Milford et al. 2004; Widloski et al. 2018; Hwang et al. 2024), including the problem of learning by combining sequential observations into a coherent map. Recent models have made significant progress in elucidating how the hippocampal formation might acquire and store knowledge about the relational structure of explored spaces (Whittington et al. 2020; Raju et al. 2024; Sharma et al. 2022; Chandra et al. 2025; Stachenfeld et al. 2017), across spatial and non-spatial domains (Killian et al. 2012; Constantinescu et al. 2016; Aronov et al. 2017; Neupane et al. 2024). Moreover, recent work links the large capacity of an invariant grid cell code (Fiete et al. 2008; Sreenivasan and Fiete 2011) to the ability to form a large number of non-overlapping maps and ameliorate catastrophic forgetting in sequential memory acquisition (Kymn et al. 2024; Chandra et al. 2025; Fiete et al. 2008). Despite this progress, it remains unknown how the hippocampal formation interacts with cortical circuits to translate these maps into goal-directed behavior and generalize across different operant tasks and conditions.

To probe relational generalization and its neural basis, we designed an abstract sequential image navigation task in which subjects mentally navigate between start and target images drawn from a fixed sequence. The task affords a testbed for a number of different generalizations. First, **mnemonic generalization**: subjects transition from visually-guided navigation, with online visual feedback, to mental navigation, where feedback is removed and the trajectory through image space must be traversed internally. Second, **transitive generalization**: subjects navigate between previously unpracticed start–target pairings, precluding recall of past traversals. Third, **structural generalization**: the abstract task structure is transferred to new environments composed of different image sequences. Additionally, we studied **avoidance of catastrophic forgetting**, a phenomenon that artificial neural networks exhibit, by testing performance in the first environment during and after the learning of new environments. We tested humans on these behavioral hallmarks of generalization (and monkeys on a subset of them), built multi-region circuit models grounded in entorhinal–hippocampal theory to identify candidate mechanisms underlying each form of generalization, and recorded from the entorhinal and parietal cortices of monkeys to validate and refine models.

We report that humans and monkeys successfully performed all forms of tested generalization, while conventional neural network models (feedforward neural network, recurrent neural networks) failed at all of them. By contrast, a modular circuit model – in which a structured entorhinal-hippocampal memory circuit, Vector-HaSH (Chandra et al. 2025) is coupled with a policy-learning network – reproduces the full pattern of human and monkey behavior. It exhibits mnemonic, transitive, and structural generalization without catastrophic forgetting. Matching monkey and human generalization behavior is non-trivial, and the model achieves it through a central principle of **factorization**: it separates *what* is being navigated (the content of an image sequence, stored as associative memory) from *how* navigation unfolds (a low-dimensional dynamics over abstract location, driven by self-generated velocity signals). Because the navigation dynamics are content-independent, the policy learned in one environment transfers immediately to new environments, enabling rapid learning without overwriting old memories.

This architecture makes specific predictions about how computation is distributed across regions, which we test against electrophysiological recordings from the entorhinal cortex (EC) and posterior parietal cortex (PPC) in monkeys. The neural data motivated a refinement in which the action policy network itself splits into two subnetworks with distinct dynamical signatures – one more strongly tracking distance to target, the other more strongly controlling action (start in the right direction and stop). The model predicts, and the recordings verify the prediction, that EC and PPC populations differ systematically in how they support the overall task-relevant computations. Finally, we show that aligning the model’s internal velocity gain with the spacing of the image sequence is required for optimal performance, predicting a gradual learning process that calibrates self-motion signals to external structure - consistent with known velocity-and gain-modulating cell types in the entorhinal–hippocampal circuit and associated cortical areas (Iwase et al. 2020; Hardcastle et al. 2017; Whitlock et al. 2008, 2012; Alexander and Nitz 2015; Alexander et al. 2023; Dannenberg et al. 2019). Overall, our work elucidates how the brain leverages memory and past learning to support data-efficient, generalizable new learning and behavior without forgetting.

## Results

We designed an abstract sequential image navigation task to probe relational generalization in humans and monkeys. Subjects navigated from a randomly selected start image, drawn from a fixed linear sequence of images (Fig. 1a), to a designated target image elsewhere in the sequence. Movement through the sequence was controlled by deflecting a manipulandum left or right, producing motion at constant speed; subjects stopped by releasing the manipulandum (Fig. 1b and Methods). We collected behavioral data from seven human subjects who completed six sessions each. In a given 1-hour session, subjects completed on average 470 trials.

**Fig. 1.**
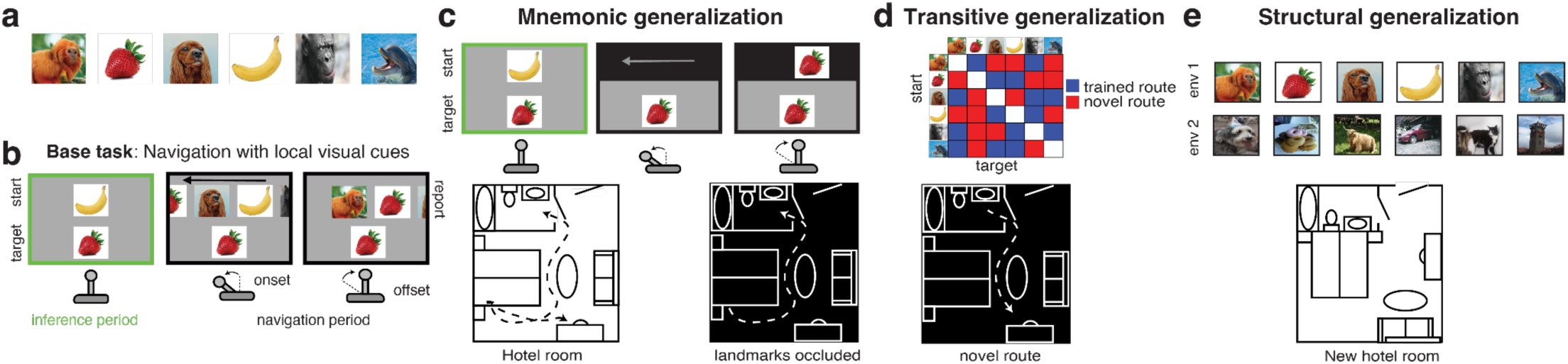
Abstract sequential image navigation task involving different types of generalization. **a.** The image sequence used for the navigation task. **b.** Base training task. Visual navigation (vnav) task in which agents are initially given a start and target image pair (left, inference period), then can deflect a joystick (left or right) to move through the image sequence at a constant speed, with visual feedback in the form of scrolling images. Agents report reaching the target by returning the joystick to the neutral position. **c.** Mnemonic generalization task. Same as (b) with no visual feedback about current location until after the joystick returns to the neutral position; i.e., mental navigation (mnav). Hotel room example illustrating mnemonic generalization when lights are off (dark background). **d.** Transitive generalization task. Same as (c), but on previously unexperienced routes (novel start-image pairings). Blue: start-target image pairings used in training. Red: novel pairings. Hotel room example illustrating transitive generalization when taking a new route in the dark. **e.** Structural generalization task. Same as (c), but with a new set of images; i.e., new environment. Hotel room example illustrating structural generalization when navigating a room with a different layout.

In the initial phase (visual navigation, vnav), subjects received continuous visual feedback showing the current image and its immediate neighbors. Because the full sequence was never simultaneously visible, successful performance required learning the relational order of images – that is, building a cognitive map of the abstract image space. Subjects then transitioned to mental navigation (mnav), in which images were occluded during movement (Fig. 1c). mnav requires internally simulating one’s current location in the cognitive map by integrating self-generated movement signals against the learned inter-image structure – the **mnemonic generalization**. To probe **transitive generalization**, we tested subjects on novel start–target pairings not encountered during training, which precludes recall of previously executed trajectories (Fig. 1d). To probe **structural generalization**, we trained human subjects on new image sequences (new environments), testing whether the abstract task structure transferred independently of sequence content (Fig. 1e). Performance on the original environment was probed intermittently throughout, allowing us to assess catastrophic forgetting.

### Mnemonic Generalization (same environment, no visual feedback)

All subjects gradually acquired vnav, reaching high accuracy within the first 10 mins (4-8 epochs; Fig. 2a, left). Upon switching to mnav, performance was immediately high (Fig. 2a, right): accuracy in the first mnav epoch was significantly greater than in the first vnav epoch, and nearly matched accuracy in the final vnav epoch (Fig. 2a, left and middle). We further quantified performance by regressing estimates against true start-target distance (Fig. 2b, left and middle) and comparing regression slopes across conditions (Fig. 2b, right). Together, these results indicate near-zero-shot transfer from vnav to mnav.

**Fig. 2.**
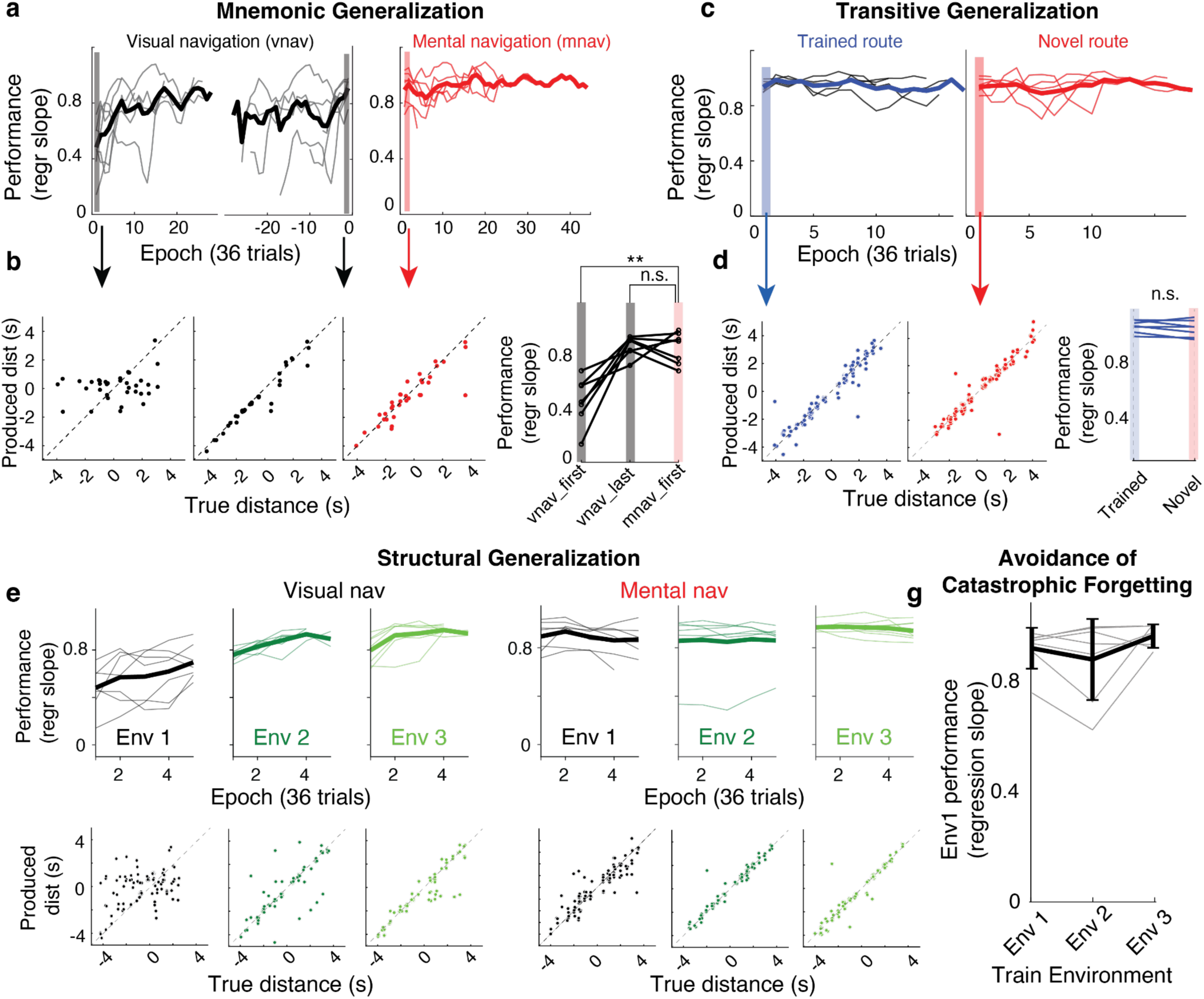
Human subjects efficiently generalize and sequentially learn new environments without forgetting. **a.** Mnemonic generalization. The performance of individual subjects on visual navigation (vnav) (left: aligned to day start, middle: aligned to vnav-to-mnav transition) and mental navigation (mnav, right) across trials within the first day. Each data point represents a regression slope relating the produced temporal distances to the true temporal distances within a 36-trial running window. Thin lines represent individual subjects, and the thick line represents the average across subjects with attrition. Vertical shades represent the corresponding epochs used to statistically compare the initial performance in vnav to that in mnav. **b.** The performance of one subject on the first epoch vnav (leftmost, black dots), on the last epoch of vnav (middle left, black dots) and on the first epoch of mnav (middle right, red dots) later in the same session. Each data point represents the true distance (measured in seconds as temporal distance) on a single trial plotted against true distance. Right plot: Comparison of performance across all subjects (regression slope) on the first epoch of vnav and that of mnav shows significant difference (two sample t-test, t(12) =-4.83, pval=0.0004), while the last epoch of vnav and the first epoch of mnav do not (t(12) = 0.3, pval=0.8). **c.** Transitive generalization. Performance of individual subjects on mnav task for comparing learning performance on familiar (blue: trained start, target pairings, left) and new (red: test start, target pairings, i.e., generalization pairs, right) routes. Data is plotted similarly to (a). **d.** The performance of one subject on mnav in familiar route (left) and new route (middle). Right: No statistical difference in performance between familiar and novel routes (two sample t-test, t(12)=0.60, pval=0.56). **e.** Structural generalization. Regression slope on the vnav task, comparing learning speeds across novel environments. Thin lines represent individual subjects; the thick line denotes the cross-subject mean. Distinct shades of green indicate Environments 1, 2, and 3. Bottom: Produced versus true distance for a representative subject during the first epoch (36 trials) of the vnav task across all three environments. **f.** Same as (e) for mnav task. **g.** Avoidance of catastrophic forgetting. Performance of all subjects, tested on the first environment (y-axis) as they learn new environments (x-axis). Thick lines represent the average across subjects.

### Transitive generalization to novel routes (new start-target pairs)

To test whether this transfer ability was supported by traversal through an internal cognitive map of the image space, we then tested generalization to new routes (Fig. 2c-d) without additional experience or visual feedback. Performance on novel routes was as good as performance on familiar routes (Fig. 2c-d), similar to non-human primates (Neupane et al. 2024). This result suggests that subjects learned and used structural knowledge (the cognitive map) of the environment to enable zero-shot inference along novel routes in a familiar environment.

### Structural generalization (new environment, same task structure)

We compared learning curves across the three sequentially trained environments. Fig. 2e-f (bottom) show one subject’s performance over the first 36 trials (1 epoch) in each environment for vnav (Fig. 2e, bottom) and mnav (Fig. 2f, bottom): performance during the early trials in the second and third environments was markedly better than in the first. The same pattern held across subjects (Fig. 2e-f, top), with faster acquisition and higher asymptotic performance in new environments, indicating transfer of structural knowledge.

### Avoidance of catastrophic forgetting

To test whether subjects exhibit catastrophic forgetting of old learning as they learn to navigate new environments, we inserted catch epochs in which subjects were returned to the first environment while learning new ones. Despite increased memory load across the second and third environments, subjects’ performance on the first environment remained undiminished (Fig. 2g).

Overall, the range of behavioral generalizations we observed in the image navigation paradigm strongly support the idea that humans build and use cognitive maps of abstract traversed spaces to facilitate highly flexible transfer and generalization and to mitigate catastrophic forgetting. This quantification of performance across generalizations also provides a human baseline for our neural circuit models of memory and generalization.

### Entorhinal-hippocampal memory-augmented action policy model

We built a multi-region brain model, which we refer to as the Vector-HaSH-augmented Action policy (VHA, Fig. 3a), with two main functional components. The first is an entorhinal–hippocampal memory circuit based on Vector-HaSH (Chandra et al. 2025), in which sensory inputs are bound - one-shot, via Hebbian plasticity - to internal scaffold states formed by the joint activity of multi-period grid modules and a hippocampal layer (Mocle et al. 2024; Hafting et al. 2005; Stensola et al. 2012). As the agent moves, velocity input updates the grid phases, advancing the scaffold state through the map. The result is a structured cognitive map that decouples *what* is at each location (sensory content, rapidly learned) from *where* one is in the map (internal dynamics, fixed across environments). The second component is an action policy network, modeled as an RNN corresponding to the role played by entorhinal and parietal cortices, which instruct actions (Fig. 3a).

**Fig. 3.**
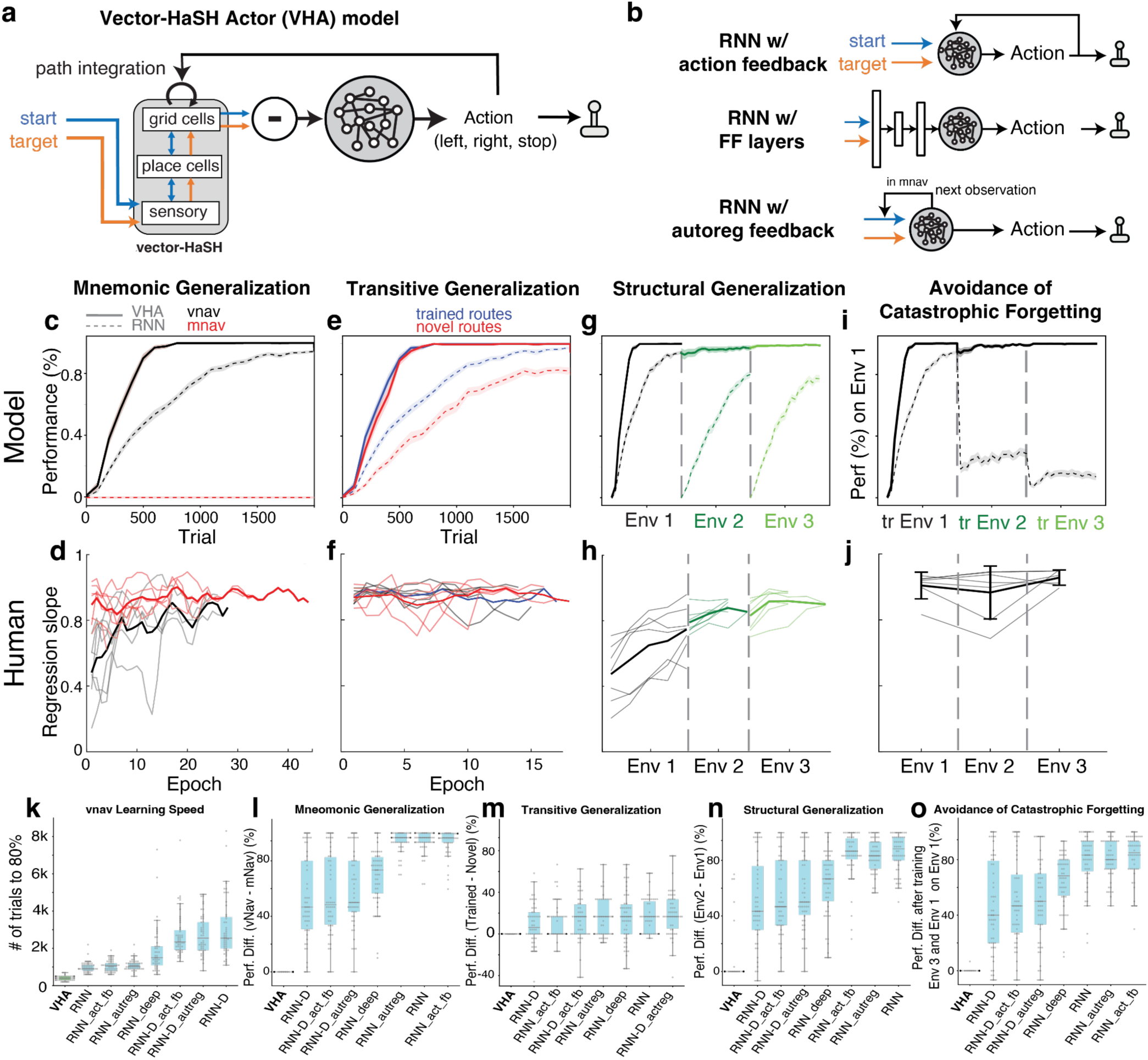
The VHA model exhibits generalizations of all three types, resembling human performance, while RNNs fail. **a.** Schematic of the Vector-HaSH-augmented Actor network (VHA) model. The current and target states are separately encoded into grid cell activations using Vector-HaSH (Chandra et al. 2025). The difference between these two states is the input to the action RNN network. **b.** Architectures of conventional RNN model variants without the VHA memory scaffold network (see Fig. S1 for full details). **c.** Model mnemonic generalization: Performance on the visual navigation (vnav) task (black), where all images are visible, and on the mental navigation (mnav) task (red), where all intermediate images are occluded. Solid lines denote the performance of the VHA model and dashed lines denote that of the conventional RNN, which fails to learn mnav without specific training on mnav. Both models were trained on vnav and tested on mnav. **d.** Same as (c) for human subjects. Thin lines represent individual subjects and thick lines represent the average across subjects with attrition. **e.** Model transitive generalization: Performance on trained routes (blue) and novel routes (red) during mnav. **f.** Same as (e) for human subjects during mnav. **g.** Model structural generalization: Performance on vnav in three different environments (black, green, light green, respectively). **h.** Same as (g) for human experiment during mnav. **i.** Model performance on vnav in the first environment (env 1) while training on other environments. **j.** Same as (i) for human experiment during mnav. **k.** Number of trials required to reach 80% accuracy for different models. **l-n**. Performance of different models for mnemonic (l, vnav vs. mnav), transitive (m, trained vs. novel routes), and structural (n, end of environment 1 vs. onset of environment 2) generalization. **o.** Performance of different models in environment 1 after training on environment 3.

When a start or target image is presented, it activates the associated grid–hippocampal scaffold states via the previously bound sensory inputs; the difference between these two grid states encodes a distance and direction in an abstract, content-independent space. The action policy network reads out this difference vector to produce a sequence of left/right/stop actions. Critically, each action is also fed back as velocity input to the grid layer, advancing the scaffold state internally – so the agent can simulate its trajectory through the image space without further sensory input. We hypothesize that this architecture will support all three generalizations from a single training regime: distance/direction extraction from grid-state differences enables traversing novel routes (transitive generalization), velocity feedback into grid cells enables mental navigation (mnemonic generalization), and content-independent grid dynamics enable transfer to new environments (structural generalization). We trained the model on the visual navigation task using a subset of start–target pairs and tested the three forms of generalization, along with susceptibility to catastrophic forgetting (Fig. 1c-e). We compared the full model with a conventional RNN and variants of VHA in which one or more of the modules were excluded (Fig. 3b, S1).

### Model behavior

The VHA learned the visual navigation task approximately three times faster than a conventional RNN (Fig. 3c, solid vs. dashed black lines). To test mnemonic generalization, we ran probe trials throughout vnav learning, with sensory feedback removed. The VHA transferred immediately without performance loss at every stage of vnav training (Fig. 3c, red solid line, fully overlapping the black solid line), matching the near-zero-shot transfer observed in human subjects (Fig. 2b, right). The baseline RNN, by contrast, continued to improve on vnav but failed to generalize to mnav at any point during training (Fig. 3c, dashed vs. solid red lines). Thus, while both models can learn to navigate with continuously available visual cues, only the VHA architecture supports navigation when those cues are removed.

To test whether the VHA solves the task by retrieving a map of the environment and inferring its location within it, we measured transitive generalization: performance on novel start-target pairings during mnav, in the absence of visual input. The VHA generalized readily, with no drop in performance relative to trained pairs (Fig. 3e, blue vs. red solid lines). Combined with its vnav-to-mnav transfer, this rules out a model-free strategy of associating visual cues with action outputs. Humans (Fig. 2c-d) and monkeys (Neupane et al. 2024) show the same pattern, consistent with both biological and model agents employing a map-based strategy. Conventional RNN – testable on novel pairs only in vnav, since it failed mnav – generalized to novel routes partially and with reduced accuracy (Fig. 3e, blue vs. red dashed lines). This is consistent with one of several heuristic solutions to the task. For instance, statistical association of start images with action directions (e.g., images at the left of the image sequence typically require rightward movement) combined with online visual feedback for stopping, rather than learning of map structure, can enable moderate generalization.

To test whether the models had learned a transferable task schema, we evaluated structural generalization by training each model on three environments sequentially. Before the first trial in each environment, the model was given a single sweep through the image sequence (visual navigation, first to last image) to allow map formation; we then measured performance from the first navigation trial onward.

The VHA achieved high performance from the first trial in each new environment (Fig. 3g green and light green solid lines), mirroring the rapid generalization observed in human subjects (Fig. 2e and 3h). Conventional RNNs, by contrast, learned each environment from scratch, with a learning rate comparable to that of the first environment (Fig. 3g green and light green dashed lines). The VHA therefore learns a cognitive map and a transferable task schema; conventional RNNs learn neither.

This rapid transfer in the VHA follows from how the memory circuit handles novel inputs. When a previously unseen image is presented, it does not retrieve any existing scaffold state. The novel image triggers an intrinsic scaffold remapping dynamics to a new combination of grid module phases (Chandra et al. 2025), which – given the size of the multi-modular grid coding space – is overwhelmingly likely to be distant from previously used scaffold states (Chandra et al. 2025). The initial sweep through the new environment then binds this fresh sequence of scaffold states, generated by path integration from the already-learned action-to-velocity mapping, to the new images. Because the grid dynamics and the action-to-velocity mapping are preserved across environments, the relational structure of the new map is established immediately, and the previously learned policy applies without modification.

We tested catastrophic forgetting by probing performance on the first environment while models learned subsequent ones, as done for human subjects (Fig. 2g). Both VHA and humans maintained high performance on first-environment probes throughout training on new environments. The RNN performance, by contrast, dropped sharply on probes of the first environment as soon as a new environment was introduced (Fig. 3i, solid vs. dashed black lines). The VHA’s preservation of past learning results from each environment being bound to a distinct region of the large grid–scaffold coding space, and therefore new bindings do not overwrite old ones.

We also tested multiple augmented RNN variants as controls, including models with action feedback, autoregressive inputs, additional feedforward processing layers, a parameter-matched RNN (BigRNN) scaled to the total number of parameters of the VHA (Fig. S1). None reproduced the generalization, transfer, or memory stability observed in the VHA (Fig. 3k-o).

Together, these comparisons show that the VHA reproduces the rapid generalization, schema transfer, and absence of catastrophic forgetting observed in humans and monkeys, while none of the RNN variants do so – even when matched in capacity, augmented with action feedback, or given access to autoregressive inputs. What distinguishes the VHA is therefore not parameter count or training signal but architecture: a preconfigured, content-independent memory scaffold that decouples map structure from sensory content and supports navigation through internal velocity-driven dynamics.

### Comparison of neural dynamics in the model and the brain

Having shown that the VHA captures the behavioral signatures of human and monkey navigation, we next asked whether its internal dynamics reflect neural activity in the brain. We compared unit activations in the VHA with neural recordings from two regions in monkeys performing the same task: the entorhinal cortex (EC) and the posterior parietal cortex (PPC). Both EC and PPC are known to support mental simulation (Neupane et al. 2024; Crowe et al. 2005) and spatial reasoning tasks (Crowe et al. 2004) and they are thought to subserve complimentary computations in spatial reasoning (Whitlock et al. 2008, 2012). We therefore reasoned that activity in EC and PPC would exhibit task modulations during abstract sequential image navigation tasks and provide a benchmark to evaluate the internal dynamics of VHA.

At the single-unit level, neurons in both EC and PPC showed mixtures of periodic and ramping activity (Fig. 4a-b). Ramping activity, modulated by start–target distance, included upward, downward, and non-monotonic forms. Units in our action policy RNNs exhibited qualitatively similar response mixtures (Fig. S2a).

**Fig. 4.**
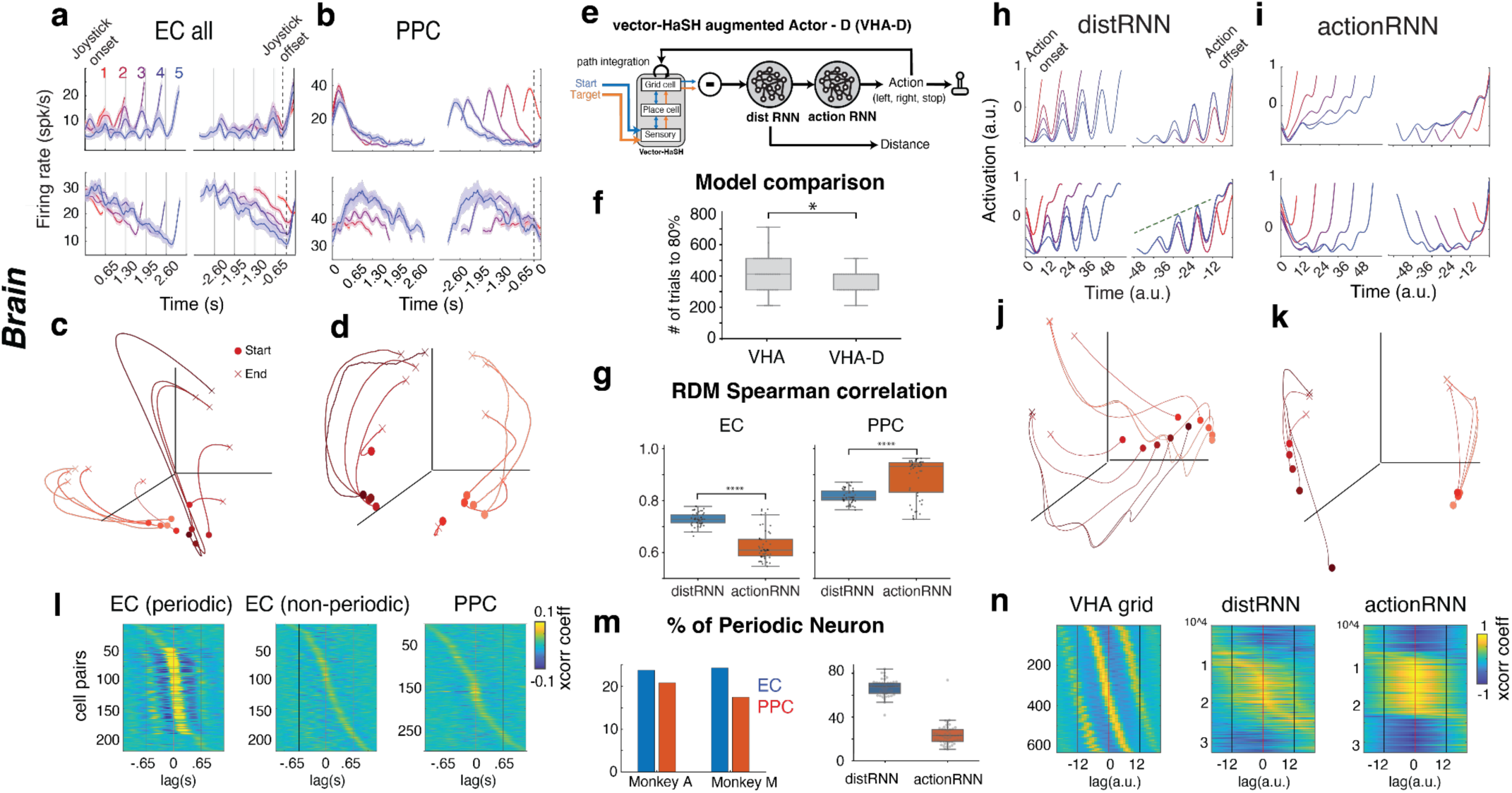
Comparison of VHA-D network dynamics with the neural dynamics in the primate brain. **a.** Firing rate activity of two neurons recorded from the primate entorhinal cortex (EC) during mental navigation. Activities are aligned to movement onset (left) and offset (right). The color gradient denotes five different temporal distances. **b.** Same as (a) for posterior parietal cortex (PPC). **c.** PCA of EC. The color gradient represents five distinct temporal distances and two directions. Trajectory start and end points are indicated by circles and crosses, respectively. **d.** Same as (c) for PPC. **e.** VHA-D architecture. **f.** Learning efficiency comparison between VHA and VHA-D. VHA-D learns faster than VHA in terms of the number of trials to achieve 80% success rate in the first environment during mental navigation. **g**. Representational dissimilarity matrix (RDM) similarity, measured with Spearman correlation between entorhinal cortex (EC) and two RNNs in VHA-D (left) and between posterior parietal cortex (PPC) with two RNNs (right). **h.** Same as (a) for distance RNN. **i.** Same as (a) for action RNN. **j.** Same as (c) for distance RNN. **k.** Same as (c) for action RNN. **l.** Population cross-correlogram heatmap of periodic neurons in EC (left), non-periodic neurons in EC (middle), PPC (right). **m.** Proportion of overall periodic units in EC and PPC of two animals (left), and distance RNN and action RNN (right) of VHA-D. EC and distance RNN are more periodic than PPC and action RNN, respectively. **n.** Same as (l) for VHA grid (left), distance RNN (middle), and action RNN (right).

At the population level, EC and PPC dynamics dissociated in a way that suggests a functional split between distance integration and action computation (Whitlock et al. 2012) (Fig. 4c-d). In EC, population responses began at a similar state across conditions and separated as the traversed distance approached the target, a signature of distance-tracking. In PPC, responses were separated early and late by *direction* with distance-graded modulation, a signature related to the action of selecting left or right manipulandum deflections. This dissociation aligns with task structure, where the first required output is a directional choice, followed by stopping, which depends on the distance traversed relative to the target. It also motivates refining the VHA model by splitting the action policy network into two functionally specialized subnetworks, which we develop next.

We split the action network into two recurrent modules: a distance network, optimized to estimate distance to the target, and an action network, which selects which direction to move (Fig. 4e). We refer to this model as VHA-D. This modular architecture improved task performance (Fig. 4f) and reproduced key features of neural activity in EC and PPC. As a comparable control, we tested task performance and learning in a model with just the two recurrent modules without the Vector-HaSH – we call this model RNN-D (Fig. S3). Neither RNN-D nor its variants reproduced the generalizations, transfer, or memory stability observed in the VHA-D.

At the single-unit level, units in the distance and action RNNs of the VHA-D qualitatively resembled those in EC and PPC, respectively (Fig. 4h-i). To quantify this, we measured each unit’s periodic modulation, computed from the autocorrelogram (Methods). In both monkeys, EC contained a substantially higher proportion of periodic neurons than PPC (Fig. 4m left). The model showed similar asymmetry: ∼70% of units in the distance RNN were periodic, compared with ∼25% in the action RNN (Fig. 4m right). Population dynamics followed the same correspondence - the distance RNN matched the convergence-then-target signature of EC, and the action RNN matched the direction-then-distance signature of PPC (Fig. 4j-k).

To quantify the similarity between neural dynamics of monkey and model, we computed the spearman correlation of representational dissimilarity matrix (RDM) (Yamins et al. 2014)(Yamins et al. 2014) (Methods) between EC/PPC, and the two policy RNNs in VHA-D (Fig. 4g). Consistent with the population dynamics, EC was more similar to distance RNN (one-tailed Wilcoxon signed-rank test, H₁: Δ_EC > 0, p<<.0001) and PPC was more similar to action RNN (H₁: Δ_PPC, p<<.0001).

The modular model makes a specific prediction about the temporal evolution of population dynamics (Narayanan 2016; Laje and Buonomano 2013; Paton and Buonomano 2018; Hardy et al. 2018; Wang et al. 2018; Soares et al. 2016; Stine and Jazayeri 2025). If EC encodes distance through path integration, its dynamics should evolve at a constant rate, independent of the total distance to the target. An action-selection network, by contrast, must transition between depress-joystick and release-joystick states regardless of the interval between start and target. As such, it should compress its dynamics over shorter intervals, producing faster trajectories for shorter distances. Indeed, trajectory speed in the distance RNN was uncorrelated or positively correlated with interval duration, while speed in the action RNN was negatively correlated (Fig. 5a left and right, respectively). Mapping the distance RNN to EC and the action RNN to PPC, the model therefore predicts a negative correlation between neural trajectory speed and temporal distance (i.e. start to target interval) in PPC and the absence of such correlation in EC. We tested this prediction in the monkey data. In one example session from EC and PPC (Fig. 5b, left and right, respectively), we found that the temporal evolution of EC population activity was not modulated by temporal distance, whereas the trajectories in PPC were faster for smaller distances, consistent with the prediction from VHA-D. However, across multiple sessions, the two distinct dynamics appeared to be mixed in the two brain areas (Fig. S4), suggesting that PPC and EC cannot be straightforwardly dissociated based on population speed dynamics. This may not be surprising given the recurrent connectivity between PPC and EC via distributed cortical-medio-temporal pathways (Amaral et al. 1983, 1987; Whitlock 2017).

**Fig. 5.**
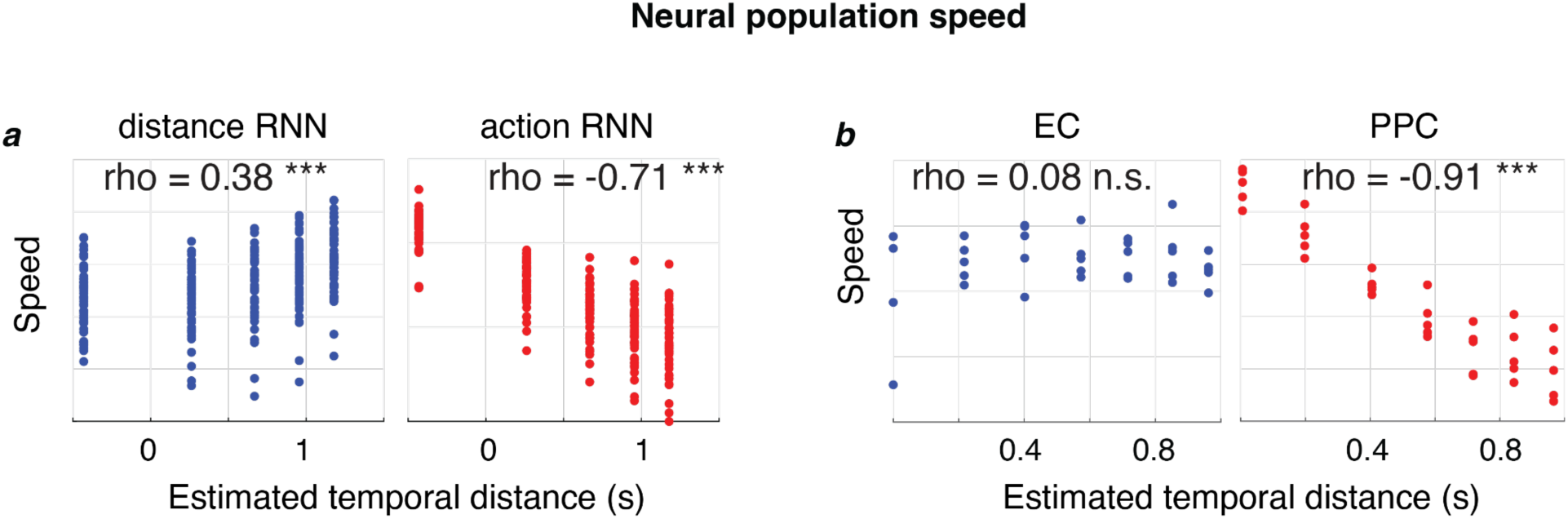
Testing VHA-D prediction of differential temporal evolution of population dynamics in EC and PPC. **a.** Scatter plot of neural speed vs temporal distance (log-log transformed) across 50 repeats for distance RNN (left, blue) and action RNN (right, red). **b.** Same as (a) for monkey EC (left, blue) and PPC (right, red).

Next, we turned to the question of the period values exhibited by the data and models. Though the image interval was arbitrarily selected, a subset of periodic units in both EC and PPC had periods that closely matched the image interval (Fig. 6a-c), with more periodic neurons at the image interval in EC than the PPC (Fig. 6c). One possibility is that the image interval coincidentally matched the period of one of the grid modules. This is unlikely, however, because only four or five discrete grid modules have been reported in EC (Stensola et al. 2012; Khona et al. 2025), and a fixed, sparse set of periods cannot be expected to contain an externally imposed interval with high probability. We therefore reasoned that the brain must actively match a module to the interval rather than select a pre-existing one, and hypothesized that the speed input to the grid-cell system is scaled so that one module’s period aligns with the image interval. Using the VHA-D model, we tested whether such a mechanism might be adaptive. First, we compared two instantiations: one in which no grid-module period matched the image interval, and one in which a single module did. Model performance was significantly higher in the matched case (Fig. 6d, Method). Next, we extended the model with a mechanism that, during task training, allows the gain of the action-derived velocity signal that drives grid-cell integration to be learned, using reinforcement learning with the negative action loss as a reward signal. Gain changes would rescale the grid period, in line with prior accounts of experience-dependent grid rescaling (Burak and Fiete 2009; Barry et al. 2007) and continuous-attractor models of velocity-gain control (Burak and Fiete 2009) (Fig. 6e; Methods).

**Fig. 6.**
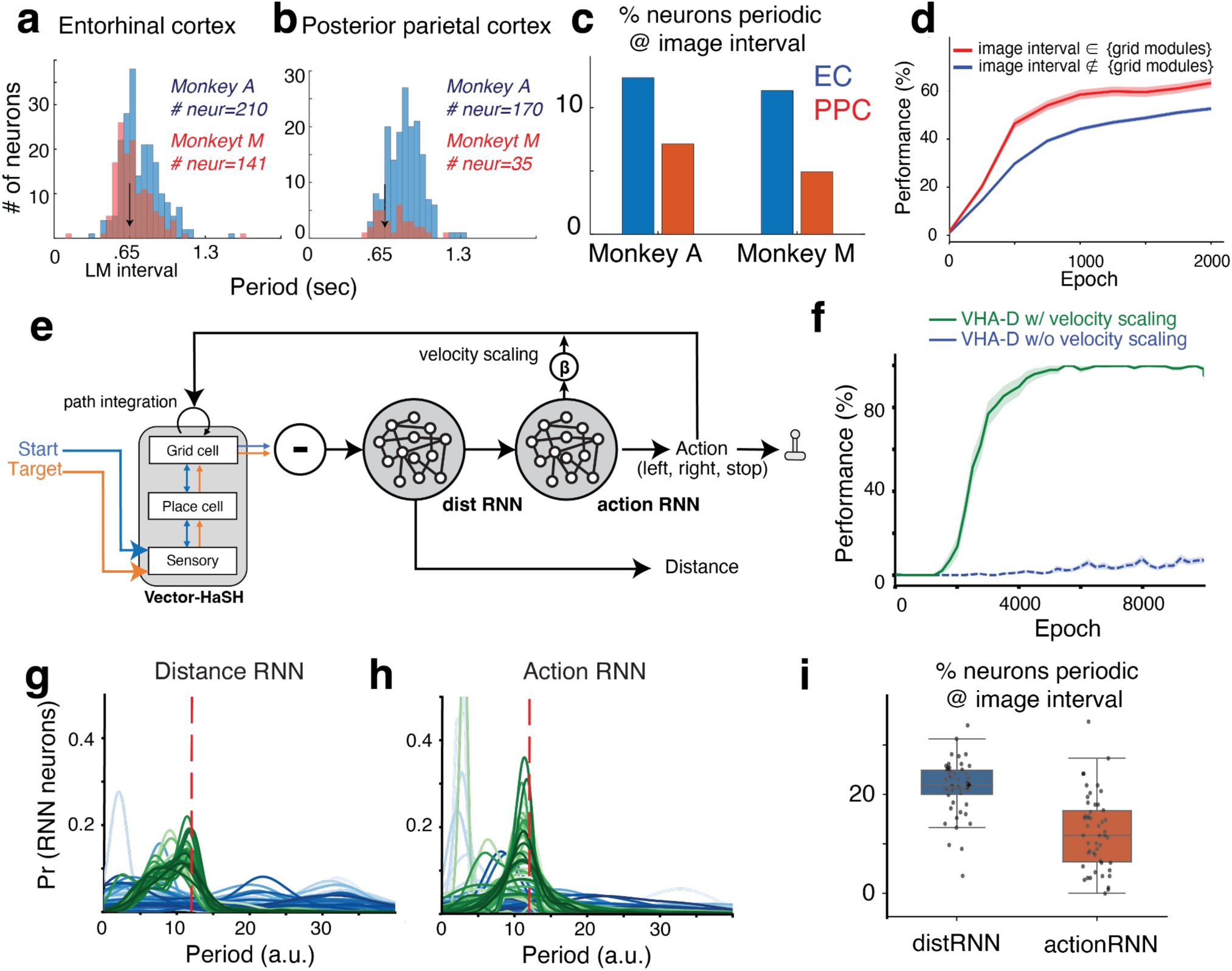
Velocity scaling leads to improved model performance and image periodicity match. **a.** Periodicity of all the periodic EC neurons in two animals. Black arrow indicates the image interval. **b.** Same as (a) for PPC. **c.** Proportion of overall periodic units whose periodicity matches the image interval in EC and PPC of two monkeys. **d.** Performance of the VHA-D model without velocity scaling in two scenarios. Red: when the image interval matches with one of the grid periodicities; blue: when the image interval doesn’t match with any grid periodicity. **e.** Schematic of the VHA-D model with velocity scaling. The action RNN predicts both the action and a velocity-scaling that scales the velocity for path integration, enabling synchronization of one of the grid periodicities with the image interval. **f.** Performance of the VHA-D model with and without the velocity scaling when the image interval doesn’t match with any grid periodicity. Velocity scaling enables the model to perform the task by matching the effective grid periodicity with the image interval. **g.** Periodicity values of neurons in the distance RNN across 50 random seeds. Each curve represents a different run. The periodicities of units in the models with velocity scaling are shown in shades of green, while those in the models without velocity scaling are shown in shades of blue. The red vertical line denotes the image interval. **h.** Same as (g) for action RNN. **i.** Proportion of overall periodic units whose periodicity matches the image interval in dist RNN (blue) and action RNN (red) in VHA-D with velocity scaling.

We tested the velocity-scaling model with a fixed image interval and many initial grid module period combinations. We found that the velocity-scaling VHA-D networks learned the task with high performance and exhibited generalization behaviors (Fig. S5b), whereas networks with mismatched periods and no velocity-scaling generally did not (Fig. 6f, blue). Then, we analyzed the periodic dynamics of our model units in both distance and action RNNs and found that the learning process resulted in one of the grid periods re-scaling to match the image interval: the periodicity distribution of trained velocity-scaling networks converged to exhibit a peak near the image period (Fig. 6g-h, green), in contrast to the networks without velocity-scaling (Fig. 6g-h, blue). Moreover, similar to monkey EC and PPC respectively, the distance RNN had more periodic neurons at the image interval than the action RNN (Fig. 6i).

Together, these results identify a mechanism by which an arbitrary external interval becomes embedded in the internal metric of the cognitive map: by adapting the velocity gain input to grid cells based on the structure of the environment, the circuit aligns one grid module’s period with the image interval. When aligned, the population state recurs consistently across images, making it straightforward for the action network to read out the location and stop precisely at the target. Without alignment, every image lands at a different combination of grid phases, leaving the downstream readout to handle a far more complex mapping. The matched periodicity observed in EC and PPC is consistent with this learned alignment.

## Discussion

Flexible behavior requires the ability to generalize past experience while preserving previously acquired knowledge. Here, we showed that humans and monkeys exhibit rich forms of generalization in an abstract navigation task and that these abilities are not captured by conventional recurrent neural networks. Instead, they are explained by a modular architecture consisting of a structured entorhinal–hippocampal memory scaffold and an action policy network.

Specifically, we found that generalization in goal-directed navigation depends on factorizing invariant structure from task-specific content. Humans and our modular network – coupling a structured entorhinal–hippocampal memory scaffold to an action policy network (VHA) – exhibited mnemonic, transitive, and structural generalization. The scaffold supplies a content-independent metric on which the policy learns to act, so that policies transfer across environments. Binding of new content to the combinatorially large scaffold states allows new environments to be acquired without overwriting old ones. Each generalization reduces, in this view, to a simple operation on the metric scaffold: transitive generalization to computing grid-state differences, mnemonic generalization to action integration over the scaffold without sensory inputs, and structural generalization to reuse of the same policy over grid-state differences regardless of image content.

A distinctive feature of the scaffold in Vector-HaSH, motivated by the invariance of grid cells across time, environments, and behavioral states (Yoon et al. 2013; Gardner et al. 2022; Trettel et al. 2019; Dong and Fiete 2024), is that its structure is innate rather than learned. This contrasts with successor representations, clone-structured cognitive graphs, and TEM (Stachenfeld et al. 2017; Raju et al. 2024; Whittington et al. 2020), which gradually learn relational structure per environment. Further, these and other existing models do not address how maps drive action and do not naturally resist forgetting. Conventional RNNs, including controls matched to the VHA in capacity, depth, and inputs, formed entangled representations of map and policy and failed at all three generalizations despite solving visual navigation on trained pairs - isolating the preconfigured metric scaffold, not parameter count or training signal, as the architectural property that does the work. VHA can be viewed as a complementary learning system with a three-way factorization – invariant scaffold, one-shot heteroassociation, transferable policy – rather than the classical two-way division into canonical fast-hippocampal versus slow-neocortical memory.

The match of the model to single-unit and population dynamics in monkey EC and PPC, in addition to the match to behavioral generalization, suggests that it captures the core mechanisms by which the brain performs these generalizations. The two-stage factorization of our action policy network, motivated by physiological differences between EC and PPC, reproduced their putative differences and made new predictions that we subsequently verified in the neural recordings: that two distinct cognitive computations are spread across EC and PPC, one in which the neural trajectory speed is uncorrelated with the start–target interval and one in where neural trajectory speed scales with it. The model’s two-stage action policy network exhibited a larger separation of these two computations than seen between EC and PPC, but the trend was directionally consistent with the regional data. It is possible that recurrent connectivity linking EC and PPC might smooth their differences, and the question of the relative role of the two regions in action policy determination is an exciting avenue for future research. Introducing recurrent connectivity between action policy modules in our model could guide neurophysiological work to investigate the mechanisms of entorhinal-parietal cortical interaction in memory guided actions.

Finally, we found that when the velocity gain driving grid integration was learnable, it converged on a value that matched grid period to image interval, reproducing the physiological finding of a match between the period of EC neurons and the image interval. The model revealed that this learned match improved task performance, proving a functional or normative explanation for the electrophysiological observation.

In sum, the hippocampal–entorhinal memory system appears to be structured to make downstream learning and computation easy. The grid scaffold converts a difficult problem – generalizing navigation across modalities, routes, and environments – into a simple one: learning a single mapping from coordinate differences to actions. The result is a system that learns fast, generalizes far, and remembers robustly, by virtue of its architecture.

## Supplementary Materials

### A Task Description

All experimental procedures conformed to the guidelines of the National Institutes of Health and were approved by the Committee of Animal Care at the Massachusetts Institute of Technology. Experiments involved seven individuals with no present or past conditions of psychiatric illness. They received a gift certificate regardless of their performance. Primate experiments involved two male, awake, behaving monkeys (species: M. mulatta; ID: A and M; weight: 8.4kg and 11.5kg; age: 6 and 11 years old). Animals were head-restrained and seated comfortably in a dark and quiet room and viewed stimuli on a 23-inch monitor (refresh rate: 60 Hz). Eye movements were registered by an infrared camera and sampled at 1kHz (Eyelink 1000, SR Research Ltd, Ontario, Canada). Hand movements were recorded using a custom single-axis potentiometer-controlled joystick, whose voltage output was sampled at 1kHz (PCIe6251 board, National Instruments, TX). We used 32- and 64-channel laminar probes (V-probe, Plexon Inc., TX) for neurophysiology recordings driven by a motorized micromanipulator (Narasighe Inc.) through a bio-compatible cranial implant. The MWorks software package (https://mworks-project.org) was used to present stimuli. Analysis of both behavioral and spiking data was performed using custom MATLAB code (Mathworks, MA).

### A.1 Visual Navigation (vnav)

Human subjects were seated comfortably in a dimly lit and quiet room and viewed stimuli on a 23-inch monitor (refresh rate: 60 Hz). We randomly selected 27 classes of images with distinctive objects from the MSCOCO dataset (Lin et al. 2014) to create a sequence containing equidistant images (6 images for monkeys and models, 9 for human subjects), denoted by *I_1_* to *I_9_*. The inter-image distance was 5 degrees of visual angle (dva). On each trial, a randomly chosen image from the sequence (*I_i_*) appears directly above the fixation point. We refer to this image as the initial image. Next, we present a different randomly selected image from the sequence (*I_j_*) directly below the initial image, which we call the target image, serving as the go cue. Subjects then press either the left or right arrow key to move the entire sequence leftward or rightward at a constant speed (10 dva/s) and stop when the image right above the target image matches it (see Fig. S6a). Trials are separated by an inter-trial interval (ITI; 750 ms). In essence, subjects must produce a 1D vector *v_p_* that matches the vector extending from the start image to the target image, denoted *v_a_*. Since the movement speed is constant, these vectors can be expressed as signed numbers whose magnitude corresponds to the temporal distance between images and whose sign represents direction. We designated rightward and leftward pointing vectors as positive and negative, respectively.

### A.2 Mental Navigation (mnav)

We used vnav to help the subjects learn the basic task contingencies, inter-image distance, image sequence, and speed. This initial training portion lasted about 10 mins for human subjects (200 trials on average +-90 trials, see Table S1) with a criterion of 75% accuracy and minimum of 72 trials to transition to mental navigation. Next, we started the training on the main mental navigation (mnav) task, which is identical to vnav, except all the images in the sequence except the initial image are invisible and remain so throughout the trial, even after the button press as the sequence starts moving covertly. After pressing a manipulandum (keyboard button for humans and joystick for monkeys), the image closest to the center of the screen is presented (Fig. 1b). On error trials, subjects were allowed to make subsequent attempts to produce a corrective vector. A cumulative accuracy value was displayed on the screen to motivate the subjects to maintain a goal of 75% accuracy. Each trial was considered a correct trial if the relative error defined as |*v_p_*-*v_a_*|/*v_a_* is smaller than a criterion value of 20% error. The rest maintained their accuracy of no less than 60%. Table S2 illustrates the sequence of tasks across the six sessions. Further experimental details of monkeys’ data can be found in (Neupane et al. 2024).

### A.3 Sessions and Curriculum

As illustrated in Fig. S7, seven human subjects completed six sessions of approximately 60 minutes each on separate days. For each image sequence, subjects initially trained on vnav trials followed by mnav trials. In the final session subjects were tested on all three sequences in both vnav and mnav conditions, with training and held-out pairs interleaved. The full session curriculum is shown in Table S2; per-subject trial counts for vnav and mnav are shown in Table S1.

### B Electrophysiology and Preprocessing

All procedures are described in Neupane et al. (2024). Briefly, all recordings were carried out after the completion of task training with a recording chamber that provided access to EC and PPC. We located EC and PPC based on stereotaxic coordinates and structural MRI scans acquired from both animals after the chamber implantation (Saleem and Logothetis 2012). To target EC reliably, we used a grid system inside the recording chamber. We registered grid holes relative to the brain using an MRI scan in which the holes were filled with an MRI contrast agent (5mg/ml Gadolinium + 10mg/ml agar). We used the registered grid system together with readings of anatomical images along the penetration path to target EC and PPC accurately.

We recorded extracellular neural activity in EC acutely across 32 sessions (A:17, M:15) and PPC across 8 (A:2, M:6) sessions using 32 or 62-channel linear V-probe array electrodes from Plexon Inc. All channels had an impedance of 275 (+-50) kOhms. Recorded signals were amplified, bandpass filtered, sampled at 30 kHz and saved using the OpenEphys data acquisition system (OpenEphys Inc., Lisbon, Portugal). We used Kilosort 2.0 software to detect and automatically sort spikes (Pachitariu et al. 2016). We used a Python-based GUI *(phy)* to verify and sort the output of the Kilosort algorithm manually. We first looked for spike artifacts that appeared in all channels and discarded them. We then looked for spikes that were unstable during a certain duration within a session. If nearby channels had clusters of spikes during those durations, we merged the two clusters of spikes if (i) they had a high correlation of spike waveform template (Pearson’s correlation > 0.9) and (ii) they were visually overlapping on the PC space computed over spike waveforms features. Next, if a given cluster of spikes clearly showed two sets of waveforms and the PC space also exhibited two clusters of clouds, we split the spikes by manually drawing a line on the PC space to separate the two clusters maximally. We included both single units and multi-units in our analyses. We considered multi-units, those clusters that had no more than two zero-crossings.

### C Neural Network Modeling

#### C.1 Simulation Setup

We model an image as a randomly generated *D*-dimensional vector considered as an encoded representation from the visual cortex. There is a 0-vector with the same size as the image presenting space between two images in monkey and human experiments. We use the inter-image interval as 12 steps following Neupane et al. (2024) that uses 650 ms (*dt*) and *τ* in the update rule of recurrent neural network (RNN) is 50 ms:

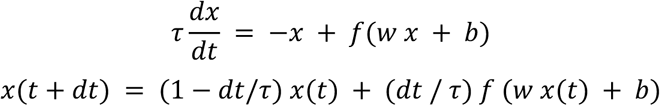

where *f* denotes activation function.

#### C.2 Vector-HaSH

Vector-HaSH is a content-addressable memory model inspired by the neural architecture of the neocortical–entorhinal–hippocampal circuit, which plays a crucial role in memory formation and retrieval in the brain (Chandra et al. 2025). In this framework, patterns are stored as stable fixed points within a neural network’s dynamics, allowing for the reconstruction of complete patterns from partial or noisy inputs – a fundamental aspect of memory recall. Vector-HaSH addresses this challenge by establishing a fixed scaffold of predefined, content-independent attractor states.

The architecture consists of three layers analogous to biological counterparts: the sensory input layer, the hippocampal cell layer, and the grid cell layer in the medial entorhinal cortex. Grid codes are employed as labels instead of the *k*-hot labels used in previous implementations. The hippocampal cell layer ℎ ∈ {−1, +1}*^N^*^ℎ^ represents spatial locations as binary vectors, capturing the discrete firing patterns of hippocampal cells. We verified that an alternative design of the hippocampal layer with scalar valued spatial code and ReLU activation function (Chandra et al. 2025) reproduced all the generalization behavior of the model and the brain-model comparisons of neural dynamics.

The grid cell layer *g* ∈ {0, 1}^∑^*_i_ ^λi^* is formed by concatenating one-hot vectors from multiple grid modules, each with *λ_i_* dimensions, reflecting the modular organization of entorhinal grid cells that encode spatial periodicity. The sensory input layer has a dimension of *N_s_*, corresponding to environmental cues and sensory information.

Before the model engages in learning tasks, the memory scaffold—including the states of the grid and hippocampal cells and the projections between these layers—is predefined. The projection matrix from the grid cell layer to the place cell layer, *W*_ℎg_, is randomly initialized to ensure an injective mapping, akin to the unique projections from grid cells to hippocampal cells observed in neural circuitry (Chandra et al. 2025). Conversely, the weight matrix from the hippocampal cell layer to the grid cell layer, *W*_gℎ_, is trained using a Hebbian learning rule. This rule strengthens synaptic connections based on the co-activation of neurons, thereby associating each active hippocampal cell (defining a hippocampal code) with the concurrently active grid cells (defining the corresponding grid state):

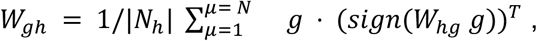

where *N* denotes the number of observed states. *μ* indexes each pattern.

As the agent navigates its environment, mimicking exploratory behavior, the weights between sensory inputs and place cells (*W_s_*_ℎ_ and *W*_ℎ*s*_) are adapted using an online pseudoinverse learning rule. This method allows for rapid, one-shot learning of associations between sensory inputs and spatial codes, consistent with the fast learning observed in the hippocampus:

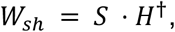

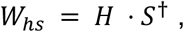

where *S* ∈ *R^Ns^*^×*N*^and *H* ∈ {−1, +1}*^Np^*^×*N*^ are matrices containing sensory and hippocampal patterns, respectively, and the dagger symbol † denotes the pseudoinverse.

The model operates by computing the activations of place cells and grid cells based on the sensory input at each time step *t*:

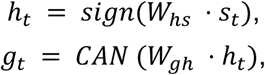

where *s_t_* is the sensory input, and *CAN*(⋅) represents the dynamics of a continuous attractor network within the grid cell layer. This network uses module-wise winner-take-all mechanisms to ensure that the grid cell activations remain valid grid states—a critical feature for accurate spatial representation and navigation.

Please refer to (Chandra et al. 2025) for more details.

The grid cell layer also integrates velocity signals (action K*a_t_* generated by RNN) to perform path integration, a process by which the agent updates its spatial representation based on movement, independent of external cues. This is achieved by shifting the activated index in each grid cell module according to the direction and magnitude of movement, reflecting the role of grid cells in encoding self-motion information:

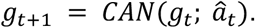

#### C.3 VHA-D: Dual-RNN Architecture

We developed a multi-region brain model to emulate the dynamics of neural systems responsible for spatial decision-making. The architecture combines two Continuous-Time Recurrent Neural Networks (RNNs) for distance and action selection with Vector-HaSH (Chandra et al. 2025), which encodes and learns associations between sensory observations and grid cell scaffolds for spatial mapping.

Given the current observation and the target image, Vector-HaSH transforms these inputs into grid states *g^curr^* and *g^term^*, respectively. These grid states represent spatial encodings akin to grid-cell firing in the entorhinal cortex.

Each grid state is then converted to an integer index per grid module, *g*′ = [*g*′_1_, *g*′_2_,…,*g*′*_K_*]. For example, if the first, third, and fifth cells are active across three modules, re, then *g*′ = [1, 3, 5]. The input to the neural network,

*z* = [*z*_1_,…,*z_K_*], is defined through modular subtraction between *g*′*^curr^* and *g*′*^term^* with respect to the grid cell periodicity *λ* for each module:

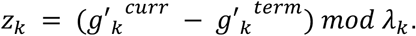

This operation preserves the periodic structure of grid-cell coding, which is important for path integration and spatial representation. The vector *z* is passed through a fully connected layer to yield *x*.

The distance RNN updates its hidden state, ℎ*^dist^*, as follows:

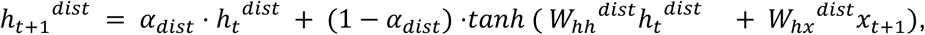

where *α_dist_* is the integration parameter (*dt*/*τ*), and *W*_ℎℎ_*^dist^* and *W*_ℎ*x*_*^dist^* are the recurrent and input weight matrices, respectively. A linear readout maps ℎ*_t_*_+1_*^dist^* to the relative distance *d*^Y^*_t_* between the current and target locations normalized by the stat-target distance.

The action RNN takes ℎ*_t_*_+1_*^dist^* as and input and updates:

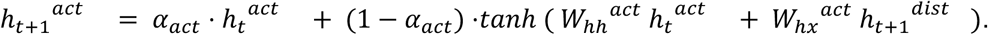

Finally, the hidden state ℎ*_t_*_+1_*^act^* is passed through two separate linear readout to predict both the action *a_t_* and the velocity scaling *β_t_*, which modulates the path integration process.

#### C.4 Velocity Scaling

We implemented velocity scaling to convert external velocity (action) into internal cognitive velocity for path integration within the grid modules of Vector-HaSH. The velocity scaling synchronizes the periodicity of the grid modules in Vector-HaSH with external periodicity (e.g., image intervals), resulting in improved performance (Fig. 6f). Given the velocity scaling at time *t*, denoted as *β_t_*, the path integration of the grid module in Vector-HaSH is computed as follows:

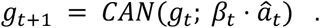

As described in the main paper, *β_t_* is predicted by the hidden states of the RNN, followed by a two-layer MLP.

### D Experimental Details

#### D.1 Objective Function

We use the mean square error (MSE) loss for relative distance, cross-entropy loss to predict the correct action of the environment and the REINFORCE (Williams 1992) to train the velocity scaling, assuming that the agents do not know their internal grid module’s periodicity.

The entire objective function, *L* for every episode, is defined as follows:

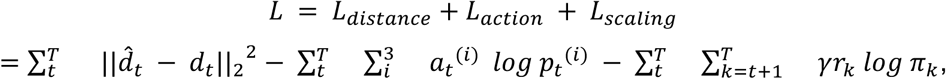

where *d_t_* and *a_t_* the ground-truth relative distance and action. *p_t_* means the predicted action probability distribution at time *t*, respectively. Similarly, *r_k_*and *π_k_* represent the reward and the probability distribution of the scale factor at time *k*. The reward is defined as negative *L_action_* at each time step and set to 100 at the final step upon successful completion of the task.

#### D.2 Details of RNN Variants

● *The RNN with action feedback*: The prediction action (scalar) is concatenated with the input of the RNN.
● *The RNN with autoregressive feedback*: A two-layer MLP is attached to the RNN to predict the next observation. This predicted observation is fed back into the network during the mental navigation period. The decoder is trained using mean squared error (MSE) loss.
● *The RNN with FF layers*: The Vector-HaSH module is replaced by a two-layer MLP with identical weight dimensions. This verifies that the parameter count (i.e., the number of neurons: 384 and 36) is not the primary factor driving performance on VHA.
● *Big RNN:* An RNN with a larger hidden size, scaled to match the total number of parameters in the combined Vector-HaSH and RNN architecture used in VHA.

#### D.3 Hyperparameter

We modeled an image as a randomly generated 384-dimensional vector (or effectively 384 pixels), and the interval as a zero vector of the same dimension. As an agent moves in one direction, the input vector is shifted by 64 pixels toward the direction, which means that each image is 6 steps and the image-to-image interval is 12 steps long. In the Vector-HaSH module, the grid periods are 11, 12, and 13 and the number of place cells is 400. The dimension of hidden states of RNN is 256 and its decaying factor *α* is 0.9 for both RNN. The hidden states pass ReLU activation followed by one fully connected layer to predict action (move left, stop moving, and move right). The relative distance is scaled up by 5. We trained the models for 2000 trials for each environment and the maximum length of each trial is 100. We use Adam (Kingma and Ba 2015) with a learning rate 0.001. For Fig. 6f, we change the second grid period to 4, 6, 8, 12, 16, 32 maintaining other periods at 11 and 13, with various image intervals (4, 6, 8, 12, 16, 24, 32). To accommodate a range of grid periods and image intervals, we changed the environment’s movement resolution. In the three resolutions we ran our model, each time step was equivalent to 1 (i.e. the original resolution) or half or one-third.

#### D.4 Evaluation Metrics

We use accuracy (average success rate over one epoch) to measure the performance of the models. The agent succeeds in an episode if it stops at the target image without changing direction and without additional stops. To measure the performance of human subjects, we calculate the regression coefficient between the produced distance and the true distance (chance performance: 0 and perfect performance: 1).

### E Quantification and Statistical Analysis

#### E.1 Firing Rate

To plot firing rates, we smoothed spike counts in 1-ms bins using a Gaussian kernel with a standard deviation of 100 ms. Because of variability in the temporal distance estimates, trials associated with the same condition (i.e., the same direction and distance) had varying lengths. Therefore, to compute trial-averaged firing rates for each condition, we used 40 ms bins for the median temporal distance estimate and appropriately stretched or compressed the bins for shorter or longer estimates, respectively (Wang et al. 2018).

#### E.2 Periodicity

We computed a periodicity index (PI) for each EC neuron using a procedure (Neupane et al. 2024) similar to that used to compute the gridness score during spatial navigation tasks. (i) We pooled firing rates for trials requiring mental navigation over at least three images to ensure that trials were long enough to compute periodicity. (ii) We truncated trials at 500 ms before the joystick offset to ensure that our estimate of periodicity was not biased by the associated large anticipatory response. (iii) We detrended firing rates using linear regression fits to the firing rate profile so that ramping activity would not mask the presence of periodicity. (iv) We computed an average autocorrelogram (ACG) for each neuron by averaging the single-trial autocorrelation function of firing rates at lags between 0 and 2400 ms. (v) To detect periodicity, we computed the correlation between the ACG and the shifted ACG for varying lags ranging from 0 to 1300 ms. (vi) We defined PI at each lag as the difference between ACG for that lag and ACG for half that lag. This procedure is analogous to how gridness scores are computed, except that instead of a two-dimensional spatial ACG, a one-dimensional temporal ACG is used. To evaluate the significance of PI for each neuron, we also created a null distribution for PI using surrogate data generated from a zero-mean Gaussian Process (GP) with a squared exponential kernel (maximum variance =1; length constant = 100 ms). To match the smoothness of GP to our smoothed firing rate, the length constant parameter of the squared exponential kernel was equal to the width of the Gaussian smoothing kernel (100ms) used for smoothing the firing rates of EC neurons. We then passed the GP surrogate data through a non-homogeneous Poisson process and smoothed the resulting spike train to obtain our surrogate GP null data. We repeated this process 1000 times to obtain a distribution. A neuron was classified as periodic if its PI at any lag was higher than 2 standard deviations from the PI obtained from surrogate data and if the total number of pooled trials was higher than 15. The trial count threshold was applied to remove spurious periodicity arising from low signal-to-noise firing rates. The results are robust to the choice of the minimum number of trials. We verified that our results and conclusions were unaffected when the trial count threshold was raised (for example, to 35 or 75). For a neuron with significant PI, its periodicity was quantified as the lag at which the PI was the maximum.

#### E.3 Neural Trajectory Speed Computation

To compute the speed of neural trajectory across temporal distances within each session, we first denoised the simultaneously recorded neural activity using Principal Component Analysis. We retained the top principal components that accounted for 80% of the variance and projected the raw activity onto them to obtain the denoised neural population activity. We then computed the total geodesic distance traversed by each trajectory and divided it by the elapsed time, yielding a measure of the neural trajectory’s speed. To increase statistical power, we uniformly divided the distribution of produced intervals across all trials into 8 time bins. We randomly subsampled a fixed number of trials from each bin and calculated the average firing rate for all neurons in each bin. We then applied PCA to this neuron x (bin x time) matrix. We repeated the sampling process with replacement to obtain 20 bootstraps of the neural speed measure. We then computed the Spearman correlation between the neural speed and temporal distance. This correlation metric indicated temporal scaling if it was significantly negative, and its gradation denoted the degree of temporal scaling (Wang et al. 2018).

#### E.4 Periodicity of Model Units

We computed the autocorrelogram (ACG) of RNN neural activity for each trial, averaged the ACGs across trials, and estimated the time lag associated with the first side lobe peak, following the method of Neupane et al. (2024). A hidden unit is considered to exhibit periodic activity if the peaks of its ACG recur at regular intervals. We denote the interval between successive peaks as the periodicity index (PI). Specifically, the autocorrelation of each hidden unit was calculated as follows:

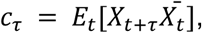

where *X*^d^*_t_* is the complex conjugate of the hidden unit’s activity, *τ* ∈ [−*T*, +*T*] represents the time lag, and *T* is the episode length. To identify peaks, we used the Python SciPy package (scipy.signal.find_peaks), which locates local maxima and applies various threshold filters. Default hyperparameters were used to detect peaks.

#### E.5 Representational Dissimilarity Analysis

We used representational dissimilarity matrices (RDMs) to compare the geometry of population activity across brain regions (EC, PPC) and the RNN modules of VHA-D (i.e, distance RNN and action RNN), following the representational similarity analysis framework (Kriegeskorte et al. 2008). Each RDM is defined over a set of task conditions that index its rows and columns; the conditions are the 10 signed image distances {-5,…,-1,1,…,5}.

For each dataset (monkey EC, monkey PPC, model distance RNN, model action RNN) and condition, we extracted the activity pattern during the navigation window: for the monkey, from joystick onset over a duration proportional to the image distance (∼650 ms per image step), using the smoothed firing rates described above; for the model, the final |d|+1 trajectory time steps, with earlier steps masked. Single-trial patterns were trial-averaged per condition, time-warped to a common length of T=12 bins and concatenated across time and units. To equalize dimensionality and suppress noise, each dataset’s per-condition trajectories were projected onto its leading 3 principal components, calculated from the concatenated trial-averaged trajectories.

We computed each RDM as the matrix of pairwise correlation distances (1-r, with r being the Pearson correlation between activity patterns across a given pair of conditions), confirming robustness to Euclidean and cosine distances. We compared RDMs by taking the upper-triangular RDM entries (excluding the diagonal) and computing their Spearman rank correlation *rho*, which is invariant to the absolute dissimilarity scale. We analyzed each brain region against the two RNN modules separately – EC against the distance and action RNNs, and PPC against the distance and action RNNs. For each brain region we asked which RNN module matched its representational geometry best. For each of the 50 model seeds we formed per-seed specificity scores Δ_EC = ρ(EC, distance RNN) - ρ(EC, action RNN) and Δ_PPC = ρ(PPC, action RNN) - ρ(PPC, distance RNN), where a positive score indicates that the region is better matched by the predicted module than by the alternative. A one-tailed Wilcoxon signed-rank test across seeds (scipy statistics toolbox, wilcoxon test, alternative=’greater’) showed EC being more similar to the distance RNN than to the action RNN (p = 1.1 x 10⁻⁸), and PPC being more similar to the action RNN than to the distance RNN (p = 9.6 x 10⁻¹⁷). We bounded the achievable similarity with a noise ceiling, estimated for each monkey region by repeatedly (100×) splitting trials per condition into two stratified halves, correlating their RDMs, and applying the Spearman–Brown correction ρ_full = 2ρ/(1+ρ).

## Author contributions

J.H. and I.R.F designed the computational experiments. S.N. and M.J. designed the monkey and experiments. J.H. conducted computational experiments and analysis. S.N. conducted monkey neural data collection and analysis. J.H. and S.N. conducted human data collection and analysis. M.J. and I.R.F. supervised the project. All authors contributed to writing the manuscript.

---

**Fig. S1.**
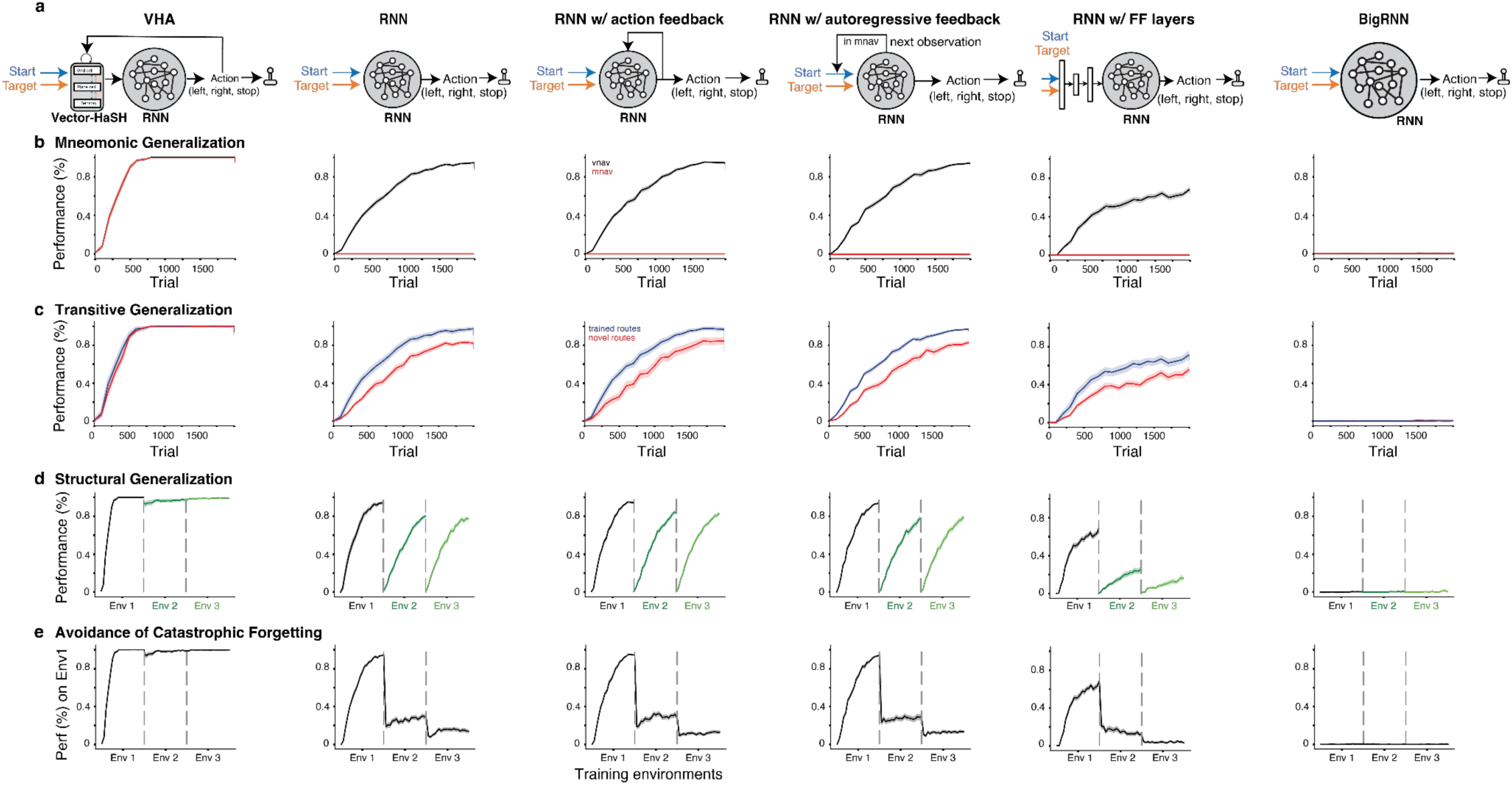
Variants of conventional single-RNN circuits fail at generalization, transfer, and continual learning. **a.** Schematic of the conventional RNN variants with four modifications: (left) with action feedback, (middle) with autoregressive feedback and (right) with feed-forward layers. **b.** Mnemonic Generalization: Performance on the visual navigation (vnav) task (black) where all images are visible and on the mental navigation (mnav) task (red) where all intermediate images are occluded.**c.** Transitive Generalization: Performance of the three models on image pairs used for training (trained routes, blue) and those held out for testing (novel routes, red). **d.** Structural Generalization: Learning performance of the models in three environments with similar structure and action dynamics but each with a unique sequence of images. **e.** Avoidance of Catastrophic Forgetting: Performance of the models in the first environment while training in subsequent environments.

**Fig. S2.**
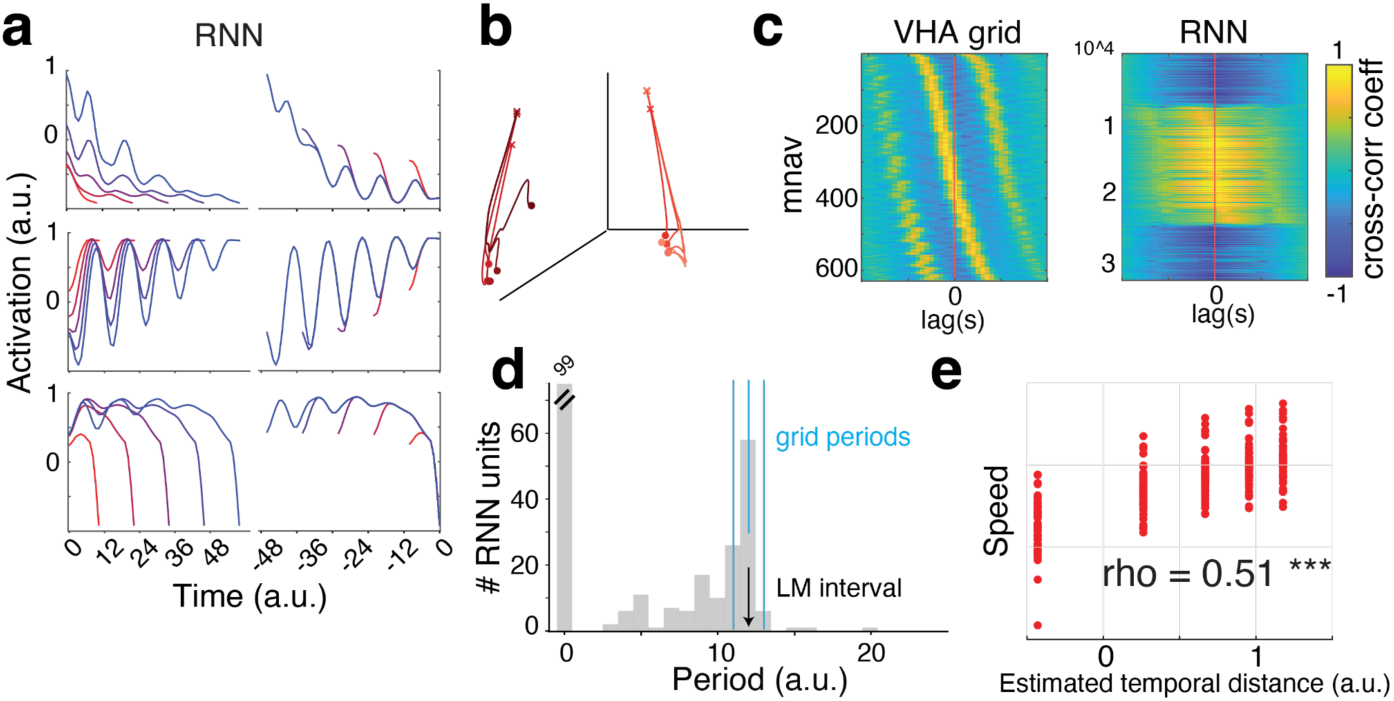
Analysis of VHA. **a.** Firing rate activity of two neurons recorded from the RNN of VHA. **b.** PCA of RNN. The color gradient represents five distinct temporal distances and two directions. Trajectory start and end points are indicated by circles and crosses, respectively. **c.** Pair-wise cross-correlogram heatmap of periodic neurons in VHA grid (left) and RNN (right), sorted by time of peak correlation. **d.** Periodicity of all the periodic neurons in the RNN of VHA. **e.** Scatter plot of neural trajectory speed (for RNN units) vs temporal distance across 50 repeats of model training.

**Fig. S3.**
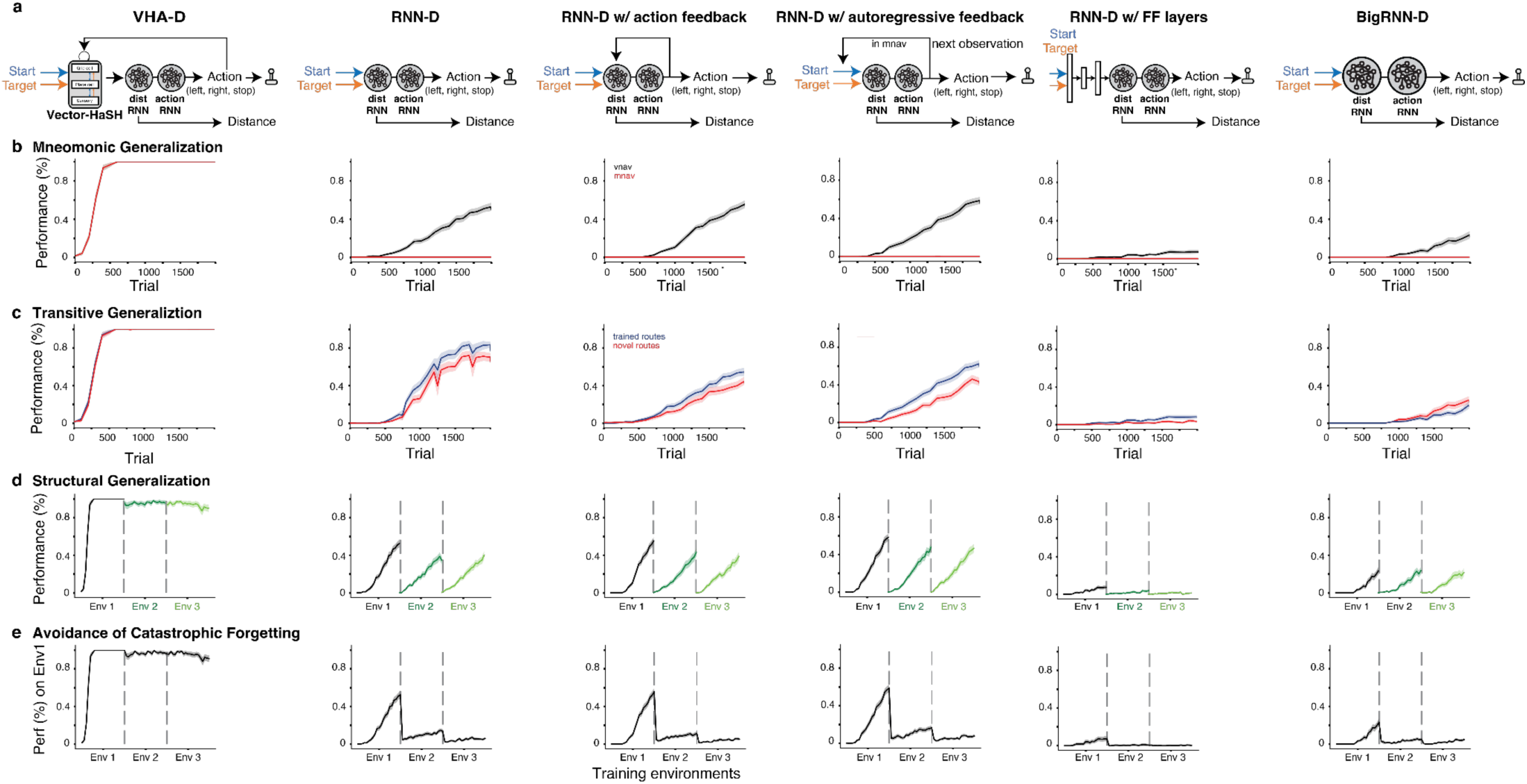
Variants of dual-RNN circuits fail at generalization, transfer, and continual learning. **a.** Schematic of the conventional dualRNN (RNN-D) variants with four modifications: (left) with action feedback, (middle) with autoregressive feedback and (right) with feed-forward layers. **b.** Mnemonic generalization: performance on the visual navigation (vnav) task (black) where all images are visible and on the mental navigation (mnav) task (red) where all intermediate images are occluded. **c.** Transitive generalization: performance of the three models on image pairs used for training (trained routes, blue) and those held out for testing (novel routes, red). **d.** Structural generalization: learning performance of the models in three environments with similar structure and action dynamics but each with a unique sequence of images. **e.** Avoidance of catastrophic forgetting: performance of the models in the first environment while training in subsequent environments.

**Fig. S4.**
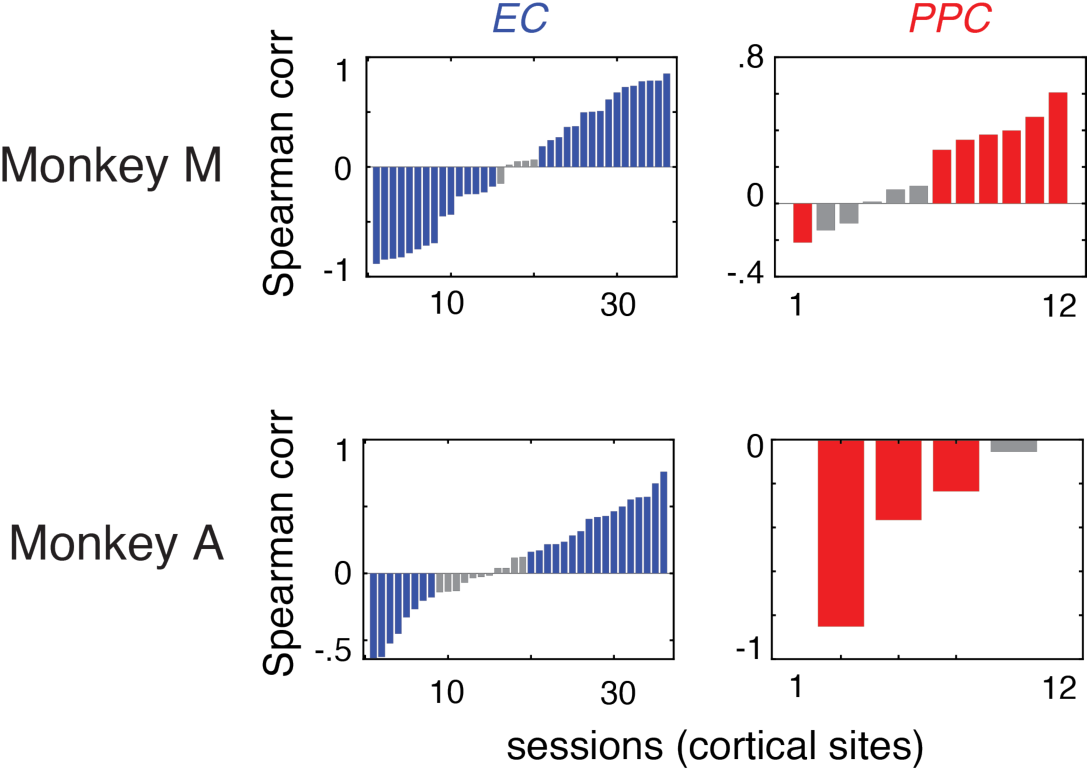
Speed control dynamics and periodicity dynamics in two animals, shown separately. Spearman correlation of neural speed vs temporal distance across different cortical sites (sessions) for EC (left) and PPC (right) for monkey M (top) and monkey A (bottom). Gray bars indicate sessions in which the correlation was not statistically significant (p>0.05). Blue and red bars denote repeats with significant correlation. Bars are sorted in ascending order of correlation values.

**Fig. S5.**
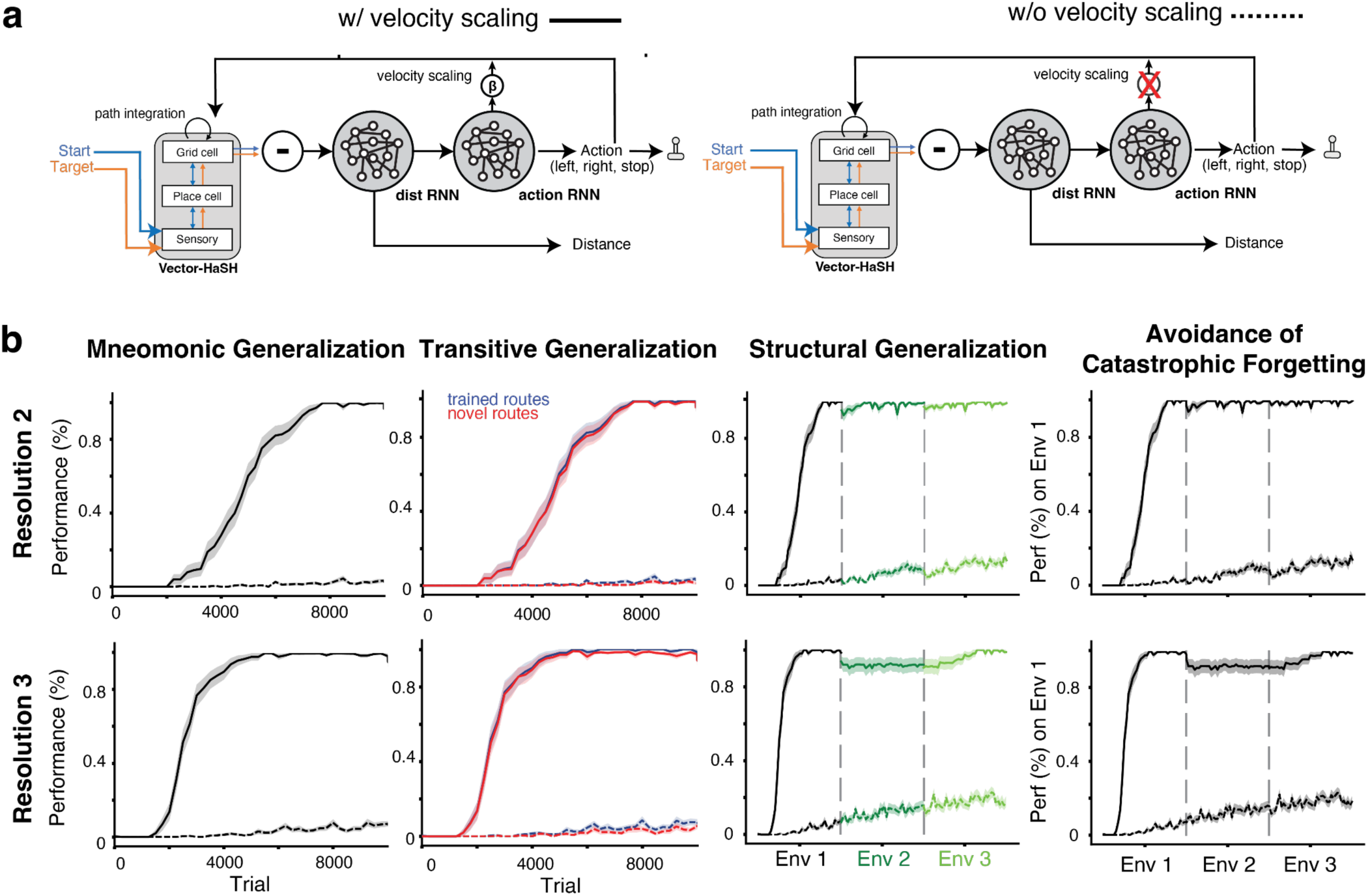
Generalization performance with and without velocity scaling in the VHA model. **a.** (left) The VHA model with an additional learning component to estimate a scale factor ‘beta’ for scaling the velocity. (right) VHA without the velocity scale factor. **b.** Behavioral performance of VHA in the mental navigation task with (solid lines) and without (dashed lines) velocity scaling on two different movement resolutions. To accommodate a range of learnable scalings, we changed the environment’s movement resolution while ensuring that our results were consistent across different resolutions (Methods).

**Fig. S6.**
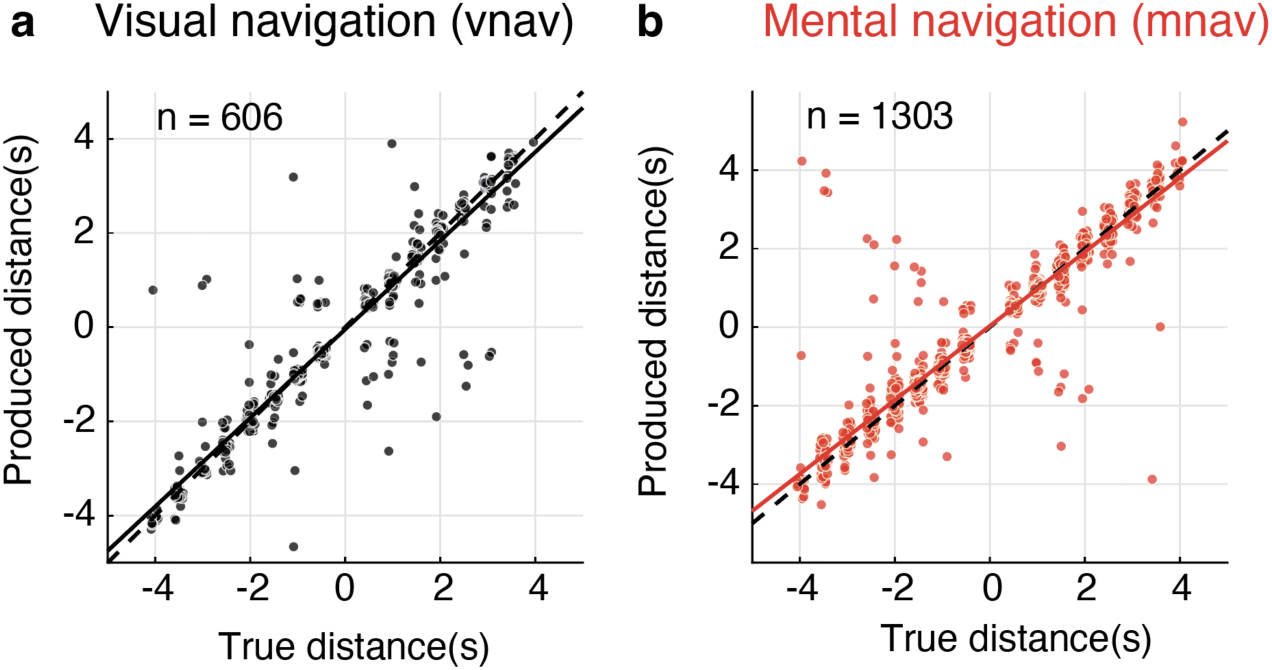
Visual and mental navigation in a single human subject. **a.** Visual navigation task performance when the image sequence drift is visible. Trials were pooled from all sessions except the first training session. **b.** Mental navigation task performance of the same subject when the image sequence images are occluded. The dashed lines in both plots represent the unity line, and the solid lines represent the regression line. Each data point represents the produced distance (measured in seconds as temporal distance) on a single trial plotted against true distance. Trials were pooled from all sessions except the first training session.

**Fig. S7.**
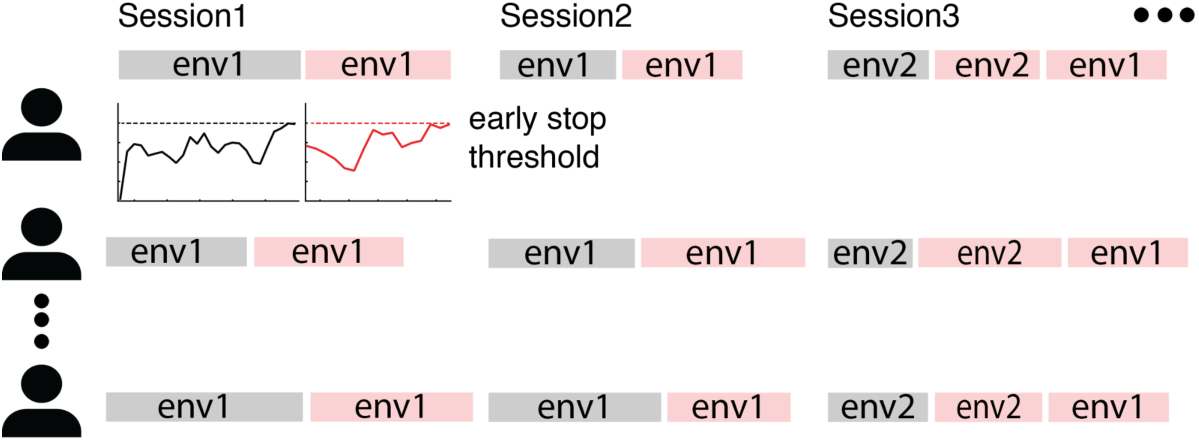
Sessions and curriculum. Seven human subjects completed six sessions of approximately 60 minutes each, on separate days. Each image sequence defines an environment (env1-env3). For each environment, subjects first trained on visual navigation (vnav) trials, followed by mental navigation (mnav) trials. In the final session, subjects were tested on all three environments in both vnav and mnav conditions, with training and held-out pairs interleaved. Per-subject trial counts for vnav and mnav are shown in Table S1; the full session curriculum is shown in Table S2.

**Table S1.**
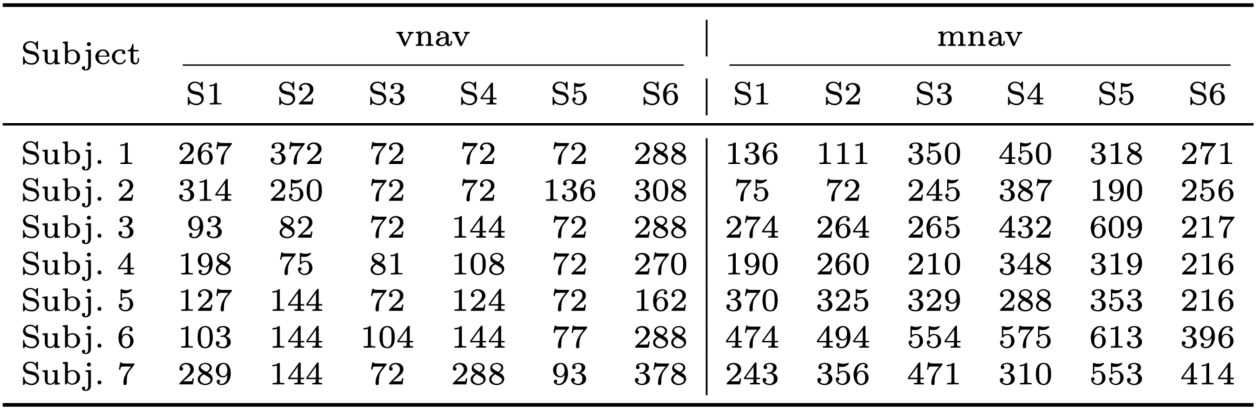
The number of trials per subject for the vnav and mnav tasks across six sessions (S1-S6).

**Table S2.**
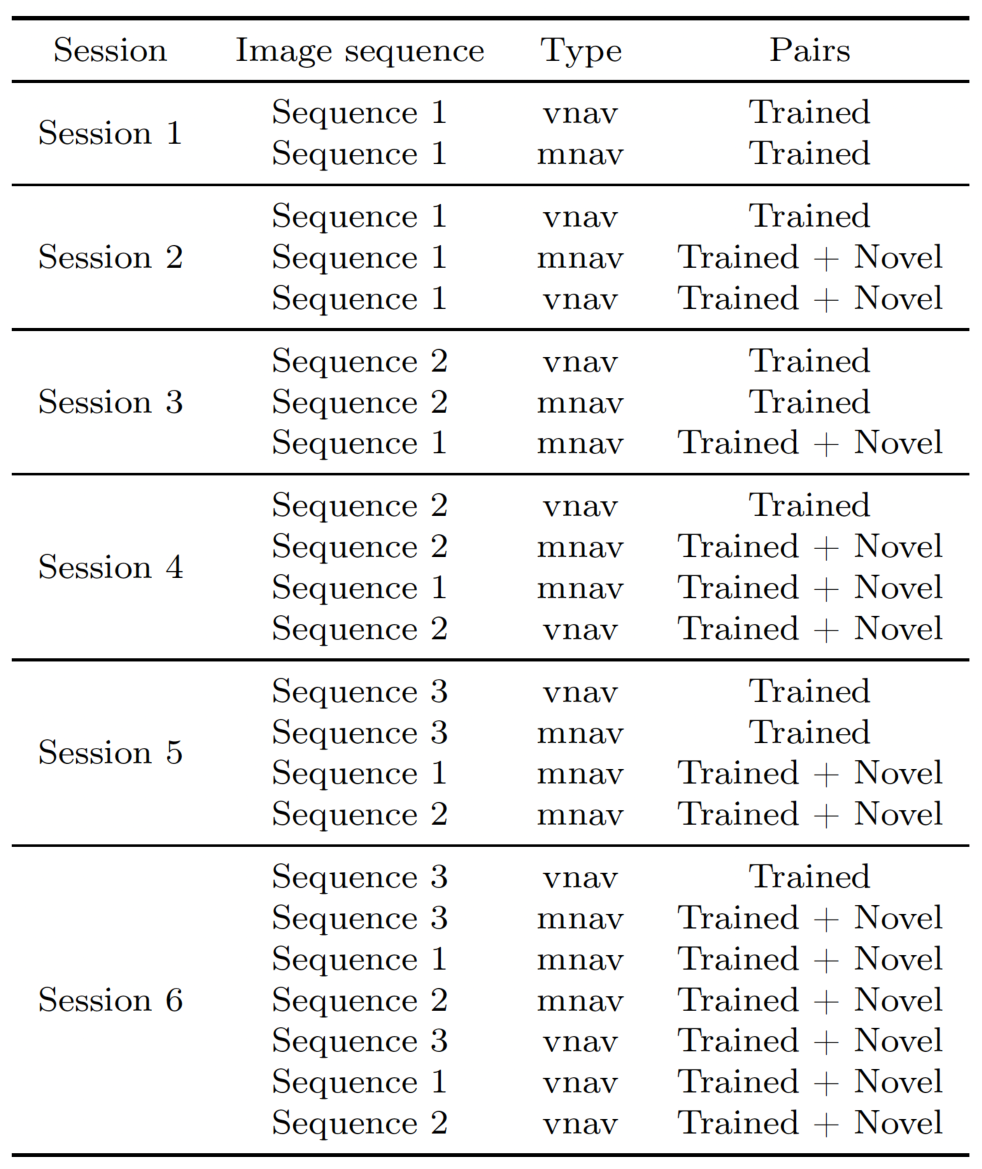
Sequence of tasks in each session for the human behavioural study. The table details the specific sessions in which each image sequence is trained as vnav and tested as mnav. “Trained” refers to the 50% of start-target pairs used during training; “Trained + Novel” indicates all pairs.

## Notes

### Competing Interest Statement

The authors have declared no competing interest.

